# Comparative single cell epigenomic analysis of gene regulatory programs in the rodent and primate neocortex

**DOI:** 10.1101/2023.04.08.536119

**Authors:** Nathan R Zemke, Ethan J Armand, Wenliang Wang, Seoyeon Lee, Jingtian Zhou, Yang Eric Li, Hanqing Liu, Wei Tian, Joseph R. Nery, Rosa G Castanon, Anna Bartlett, Julia K Osteen, Daofeng Li, Xiaoyu Zhuo, Vincent Xu, Michael Miller, Fenna M. Krienen, Qiangge Zhang, Naz Taskin, Jonathan Ting, Guoping Feng, Steven A McCarroll, Edward M Callaway, Ting Wang, M Margarita Behrens, Ed S Lein, Joseph R Ecker, Bing Ren

## Abstract

Sequence divergence of *cis-*regulatory elements drives species-specific traits, but how this manifests in the evolution of the neocortex at the molecular and cellular level remains to be elucidated. We investigated the gene regulatory programs in the primary motor cortex of human, macaque, marmoset, and mouse with single-cell multiomics assays, generating gene expression, chromatin accessibility, DNA methylome, and chromosomal conformation profiles from a total of over 180,000 cells. For each modality, we determined species-specific, divergent, and conserved gene expression and epigenetic features at multiple levels. We find that cell type-specific gene expression evolves more rapidly than broadly expressed genes and that epigenetic status at distal candidate *cis*-regulatory elements (cCREs) evolves faster than promoters. Strikingly, transposable elements (TEs) contribute to nearly 80% of the human-specific cCREs in cortical cells. Through machine learning, we develop sequence-based predictors of cCREs in different species and demonstrate that the genomic regulatory syntax is highly preserved from rodents to primates. Lastly, we show that epigenetic conservation combined with sequence similarity helps uncover functional *cis*-regulatory elements and enhances our ability to interpret genetic variants contributing to neurological disease and traits.

## Introduction

Throughout evolution, sequence divergence in the non-coding regions of the genome is believed to be a major driving force behind the emergence of species-specific traits^1^. Today, the large number of available genome sequences of eukaryotic species has allowed us to use comparative genomics to map functionally important sequences under evolutionary constraints, including *cis*-regulatory elements^2, 3^. However, sequence conservation alone cannot provide definitive evidence of the functional role of a regulatory element nor information about the cell/tissue-specificity of the elements. Additionally, not all functional elements are conserved, and some non-functional elements may appear to be conserved due to sequence similarity, leading to both false positives and false negatives in the identification of regulatory elements. Currently, we still possess a very limited base of knowledge on the evolution of gene regulatory programs. In other words, how sequence divergence leads to altered gene expression patterns across different cell types remains largely unexplored. Filling this knowledge gap is critical to understand the consequence of genetic divergence on species-specific phenotypes.

Previous bulk sequencing assays have revealed general principles concerning the conservation of *cis*-regulatory elements and tissue-specific gene expression patterns. For example, enhancers exhibit rapid turnover during mammalian evolution^4, 5^, and those conserved have lower cell type-specific activity^6, 7^. In contrast, sequence divergent enhancers play a significant role in establishing tissue and species-specific traits^8, 9^. Such divergent enhancers are often mediated by *de novo* insertion of transposable elements (TEs) carrying clusters of transcription factor binding sites^6, 10, 11^. Interestingly, the conservation of *cis*-regulatory elements^12, 13^ and expression^14^ generally decreases as development progresses. Despite the established divergence of *cis*-regulatory sequences throughout evolution, DNA motifs recognized by sequence-specific DNA binding proteins are highly conserved^15^, suggesting the existence of a conserved flexible genomic regulatory syntax. To characterize such gene regulatory syntax, it is important to perform an integrated analysis of chromatin landscape, structure, and gene expression in a cell type-specific manner across multiple species.

The primary motor cortex (M1) is a region of the neocortex preserved across eutherian mammals critical for volitional fine motor movements^16^. Recently, several reports that are part of the BRAIN Initiative Cell Census Network (BICCN) have characterized the vast complexity of cellular taxonomy, gene expression, and epigenome of brain cells in multiple mammalian species^17–21^. This included a single-cell comparative analysis of the mouse, marmoset, and human transcriptomes of M1 cells, revealing a high degree of species-specific marker gene expression^18^. However, a large knowledge gap exists in our understanding of how genome evolution influences species-specific gene expression. We, therefore, asked if sequence divergence at noncoding *cis*-regulatory elements is associated with driving species-specific traits and biology through the evolution of gene regulatory programs in different cell types.

In this study, we sought to characterize the evolution of gene regulatory programs by performing comparative epigenomic analyses Specifically, we carried out single-cell multiomics assays on brain tissue from mouse, marmoset, macaque, and humans, profiling four different molecular modalities: gene expression, chromatin accessibility, DNA methylation, and chromatin conformation. In doing so, we mapped candidate *cis*-regulatory elements (cCREs), and profiled their dynamic epigenetic states across 21 brain cell types in M1 from four species. To utilize these data for our comparative study, we developed a new framework for assessing the evolution of gene regulatory features. Our analysis demonstrates co-evolution of the epigenome and 3D genome with the transcriptome. While not all cCREs contribute to gene expression in the same way, epigenetically conserved cCREs are more likely to activate gene expression and harbor disease risk variants. Species-biased cCREs that are predicted to regulate gene expression are more likely to contribute to divergently expressed genes. Genome browser tracks are publicly available for viewing on the WashU Comparative Epigenome Browser data hub: https://epigenome.wustl.edu/BrainComparativeEpigenome/.

## Results

### Single-cell multiomic analysis of primary motor cortex in four mammals

To gain a detailed picture of how gene regulatory programs evolve, we carried out a comparative epigenomics study with the primary motor cortex of human, macaque, marmoset, and mouse (Fig. 1a). Two single-nucleus genomics assays were utilized, 10x multiome (10x Genomics) and snm3C-seq^22, 23^ (also known as single-cell Methyl-HiC), to simultaneously profile transcriptomes with chromatin accessibility and DNA methylation with 3D genome, respectively. We profiled 40,937 human nuclei, 34,773 macaque nuclei, 34,310 marmoset nuclei, and 47,404 mouse nuclei with 10x multiome and 8,198 human nuclei, 5,737 macaque nuclei, 4,999 marmoset nuclei, and 5,349 mouse nuclei with snm3C-seq (Fig. 1b). We then performed unsupervised clustering using gene expression or DNA methylation, and integrated datasets across species using orthologous genes as features (Fig. 1b, Extended Data Fig. 1a,b). Cell types were identified at subclass resolution using a combination of marker gene activity and reference mapping to the available M1 datasets from mouse, marmoset and human primary motor cortex^17, 18^. While we identified the corresponding cell types across all four species, cell type fractions were highly species-specific and changes in cell type composition correlated with evolutionary distance across species (Fig. 1c). Most notably, an expansion of the oligodendrocyte proportion and a reduction in the excitatory neuron proportion was observed from mouse to human, consistent with previous reports^24^ (Fig. 1c). Specific subclasses of excitatory and inhibitory neurons were human-enriched, such as L6 IT CAR3, Chandelier (ChC), and VIP neurons, while L5/6 NP, L5 IT, L5 ET, and PVALB neurons were consistently lower in human donors (Fig. 1d). Our data reveal trends of evolutionary divergence of cell type composition in mammalian M1, demonstrating the necessity for cell type-resolved data for cross-species comparative analysis of tissues. For downstream comparative analyses, we combined sequencing reads for each cell type, resulting in species- and cell-type resolved epigenome and transcriptome landscapes for each molecular modality (Fig. 1e).

**Fig. 1:**
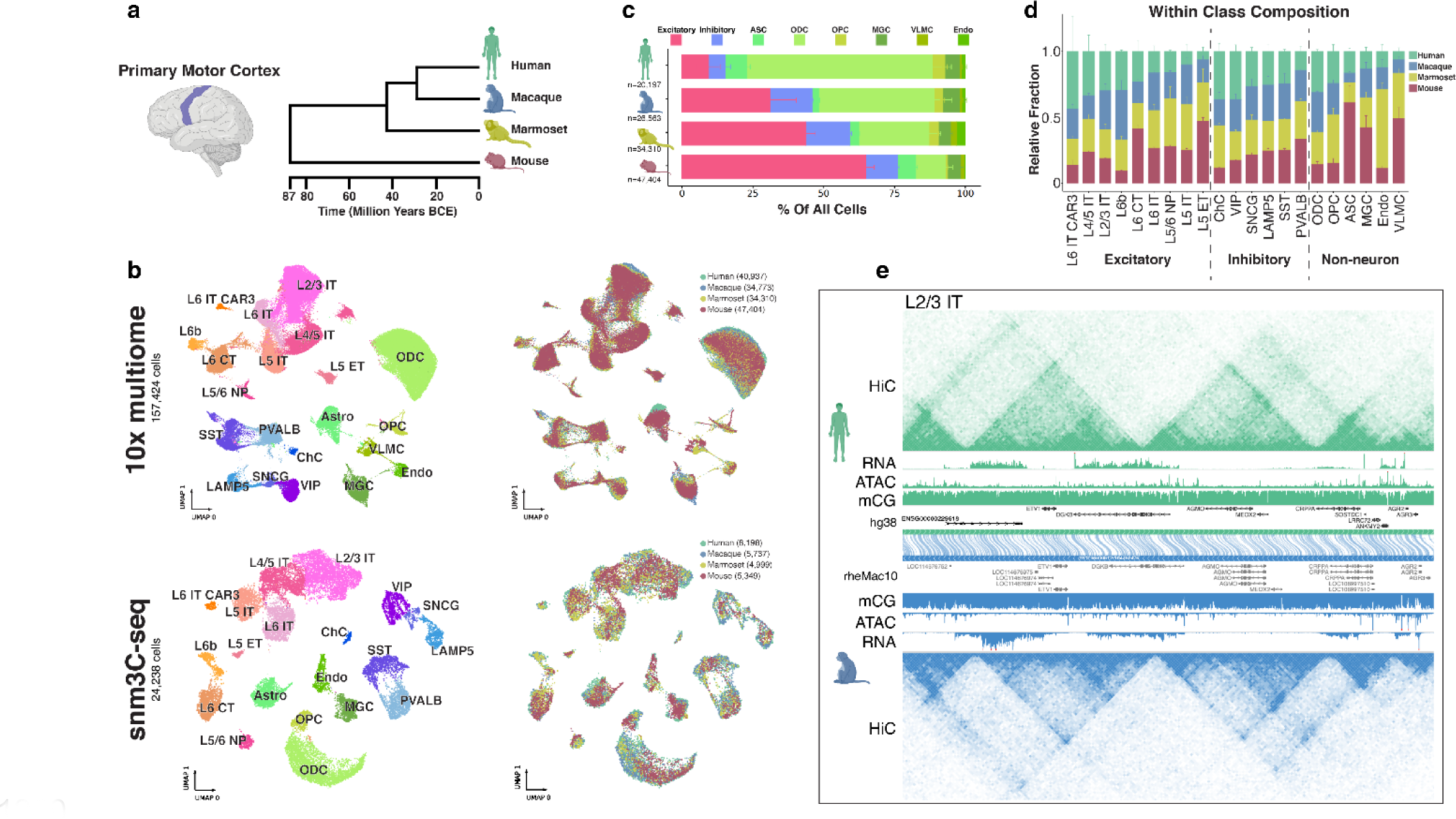
Cross-species evolutionary comparison of single-cell multiomics of the primary motor cortex. **a**, The primary motor cortex (M1) region is highlighted in an illustration of the human brain (left). Dendrogram representing evolutionary distance of each species subject to our study (right) adapted from TimeTree^51^ **b**, Uniform manifold approximation and projection (UMAP)^52^ embeddings of 10x multiome RNA and snm3C-seq DNA methylation clustering, with cell type cluster annotations (left) or species labels (right). Numbers in parentheses represent the number of cells contributed from each species. **c**, Stacked bar plots showing the fraction of each indicated cell type recovered from unbiased nuclei sorted samples. Error bars represent the standard deviation across donors (primates, n=3) or sample pools (mouse, n=8). **d**, Stacked bar plots represent the relative abundance between species of the fraction of cell type contribution to the total cells of its particular class (excitatory, inhibitory, or non-neuron). Error bars represent the same as in **c**. **e**, Screenshot of the WashU Comparative Epigenome Browser displaying a representative syntenic region alignment between human and macaque genomes with L2/3 IT data tracks for HiC, RNA, ATAC, mCG.

### Comparative analysis of gene expression across species

We evaluated the divergence and conservation of transcription between species for each gene identified as one-to-one orthologs in all four species. We defined gene expression conservation as the ability to predict the expression level of a gene in a specific cell type, given knowledge of the expression level of the same cell type in a different species. To account for the dependence relationships between cell types, we employed generalized least squares (GLS) regression^25^, for each pair of species (methods) (Fig. 2a, Extended Data Fig.2a, Supplementary Table 1). To Identify divergence of gene expression, we performed differential expression using edgeR for each cell type between each species pair^26^ (Fig. 2b, Supplementary Table 2). We identified species biased genes which are differentially upregulated in a single cell type when compared to each other species (methods).

**Fig. 2:**
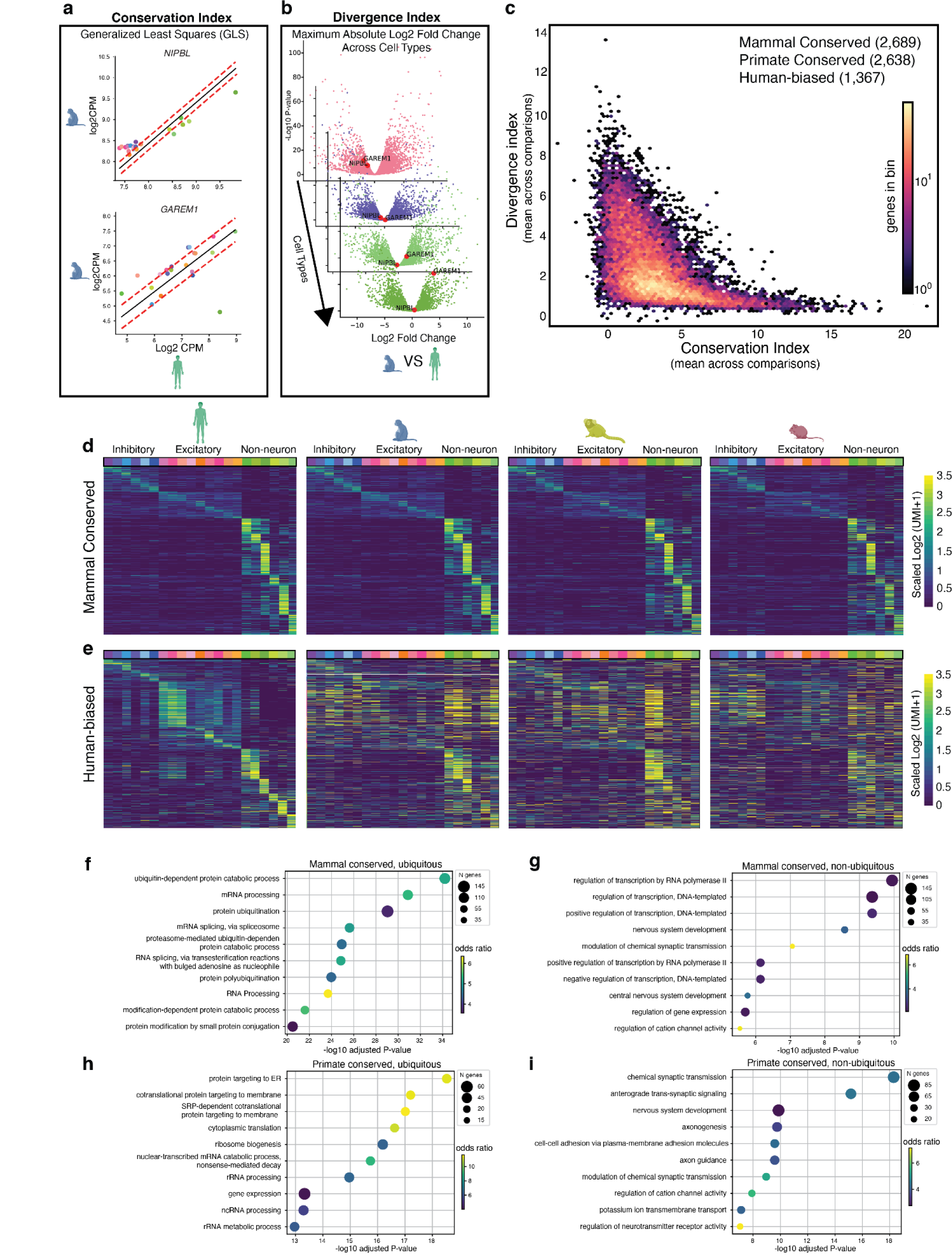
Comparative analysis of gene expression across species. **a.** Scatterplots illustrate generalized least squares regression for two genes, NIPBL (GLS T statistic= 15.460), and GAREM1 (GLS T Statistic = 3.673), between human and macaque colored by cell type. **b.** Volcano plots illustrate measurement of gene divergence as measured by edgeR between human and macaque across cell types L5IT, PVALB, Astrocytes, and Microglia. NIPBL and GAREM 1 are highlighted in red. **c.** A plot of hexagonal bins showing the relationship between average conservation index across species on the X axis, and average divergence index across species on the Y axis. The number of genes in each bin is shown by color scale. **d.** Heat maps showing patterns of non-ubiquitous mammal conserved genes across species. Gene expression is row normalized. Heat maps are ordered by the human cell type with the highest gene expression **e.** Heatmaps of human-biased gene expression patterns across species. Genes ordered by the human cell type with the highest expression. **f.** Top significant GO analysis terms for mammal ubiquitous genes. **g.** Top significant GO analysis terms for non-ubiquitous mammal conserved genes. **h.** Top terms significant GO analysis terms primate ubiquitously expressed primate conserved genes. **i.** Top significant GO analysis terms for primate non-ubiquitously expressed genes.

Of 13,822 gene orthologs across all four species meeting minimal expression criteria, we identified 2,689 (∼20%) genes with conserved patterns of expression across cell types in all species. We additionally identified 2,638 (∼20%) genes with conserved patterns of expression only among primates (Fig. 2c, Supplementary Table 3). Across species, we identify 3,511 (∼25%) genes with strong species-biased expression patterns, finding that the number of biased genes is concordant with evolutionary diversity (human: 1,376, macaque: 451, marmoset: 638, mouse: 1,367) (Supplementary Table 3).

We noted that the majority of mammal-conserved genes displayed broad gene expression patterns across cell types and so further divided them into categories of ubiquitous and non-ubiquitous expression (Extended Data Fig. 2c,d). To identify which categories of genes display high levels of conservation and divergence, we then performed Gene Ontology (GO) enrichment analysis. Ubiquitous mammal-conserved genes were most enriched in GO categories related to the regulation of protein expression, such as ubiquitin-dependent catabolic processes and mRNA processing (Fig. 2f). Non-ubiquitous mammal-conserved genes showed enrichment for such GO categories as transcriptional regulation through RNA polymerase II and DNA templated regulation, nervous system development, and cation channel regulation. Among primate conserved genes, the number of ubiquitously expressed genes dropped considerably (Extended data Fig. 5c,d), although among ubiquitously expressed genes, we saw enrichment for translational processes (Fig. 2h). Among non-ubiquitous genes, we saw strong enrichment for neuronal functions such as synaptic transmission, and axonogenesis. These differences in enrichment suggest different targets of functional conservation at different evolutionary time scales, with the stronger selection placed on genes that regulate many functions over genes encoding cell type-specific functions consistent with prior work^27^.

### Comparative analysis of chromatin accessibility across species

To identify candidate *cis*-regulatory elements (cCREs) from chromatin accessibility, we determined the open chromatin regions in each motor cortex cell type in each species with MACS2^28^ and identified 384,412 human, 336,463 macaque, 281,297 marmoset, and 333,814 mouse cCREs that display accessibility in one or more brain cell types (Extended Data Fig. 3a,b, Supplementary Tables 4-7). We leveraged liftOver^29^ to classify human cCREs as human-specific, mammalian sequence conserved (from human to mouse), or primate sequence conserved (from human to marmoset) (Fig. 3a,b, Supplementary table 8). For our comparative epigenomic analysis, we defined levels of conservation based on sequence and activity (Fig. 3a). “Species-specific” cCREs contain sequences not identified in the other three species. “Level 0” elements are cCREs with orthologous sequences across all four mammals (“mammal level 0”) or all three primates (“primate level 0”) (Fig. 3a). In addition, we performed epigenetic conservation analysis of the level 0 cCREs and defined three levels of epigenetic conservation (epi-conservation) (Fig. 3a): “level 1” are tissue conserved cCREs with a peak called across species regardless of which cell type, “level 2” are cCREs displaying accessibility in the same cell type across species, and “level 3” are cCREs with matching patterns of chromatin accessibility across all the cell types, as measured by GLS, in all the species. Analogous to species-biased genes, species-biased cCREs are defined as peaks with differential accessibility that are consistently higher in one species compared to the three other species in the same cell type.

**Fig. 3:**
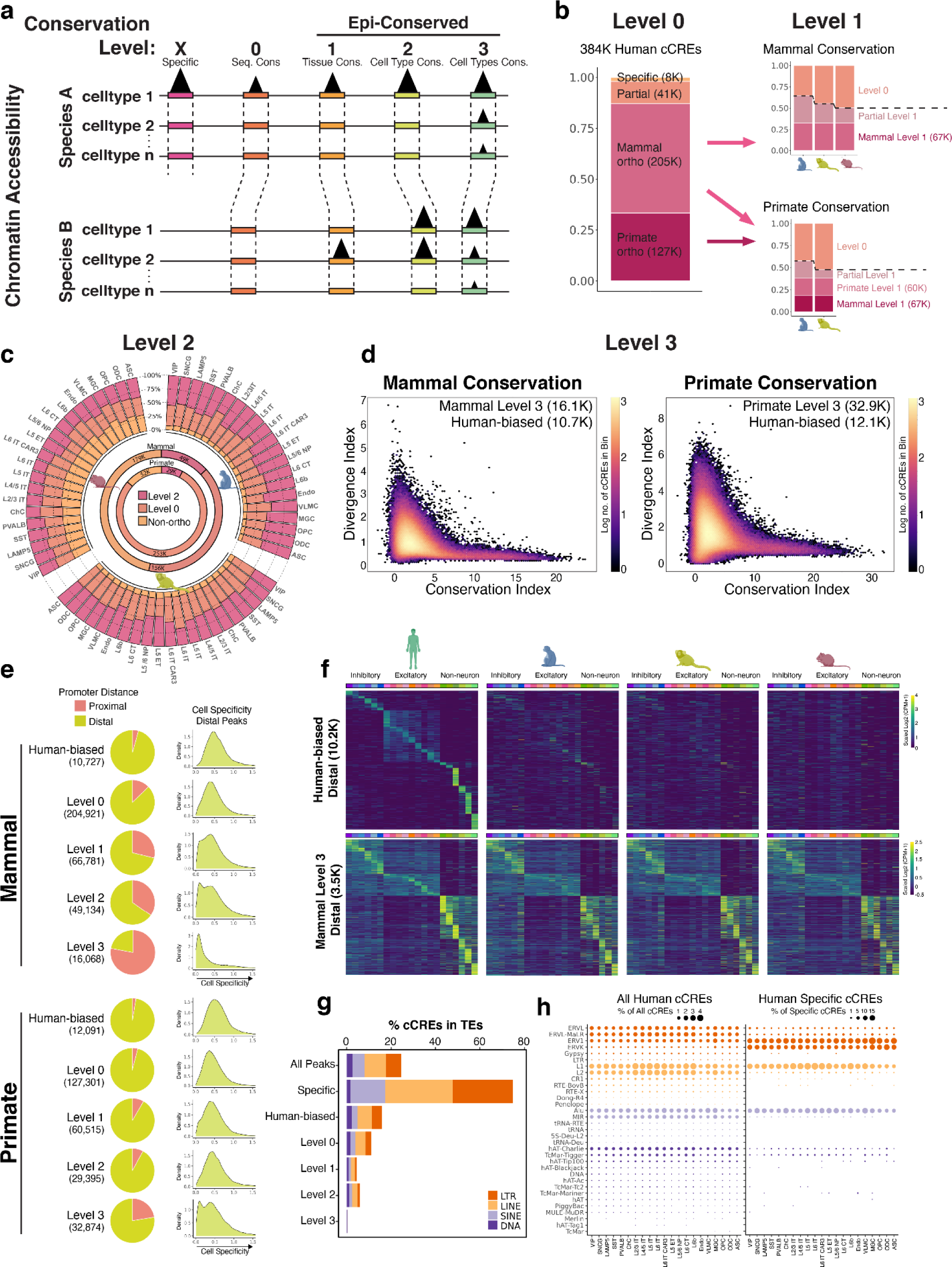
Comparative analysis of chromatin accessibility across species. **a**, Schematic illustrating levels of conservation for ATAC-seq peaks. **b**, Stacked bar plots representing the breakdown of human cCREs from ATAC-seq peaks for each indicated group for Level 0 and Level 1 conservation. **c**, Level 2 conservation of human cCREs from ATAC-seq peaks showing the overlap between each species for the same cell type (outer circle stacked bars). Inner circles show the breakdown for mammal and primate comparisons for all human ATAC-seq cCREs. **d**, Scatter plot displays the relationship between the conservation index (mean GLS T-statistic across comparisons) and divergence index (maximum absolute fold change across cell types) for Mammal Level 0 cCREs (left) or Primate Level 0 cCREs (right). **e**, Pie chart shows proportion of cCREs as promoter-proximal (≤ 1kb from a TSS) or promoter distal (> 1kb from a TSS) for indicated group (left). Density plots show distribution of cell specificity scores (methods) for cCREs in each group. **f**, Heatmaps clustered by cell type with highest signal for human-biased distal cCREs (top) and Mammal Level 3 distal cCREs (bottom). For visualization of cell type patterns of accessibility, Log2 (CPM+1) values are row scaled. **g**, Stacked bar plots showing percentage of human cCREs in TEs for different conservation groups. **h**, Dot plots showing the percentage of all (left) or human-specific (right) cCREs in different subclasses of TEs for each cell type.

Of the 384,412 human cCREs, 7,532 (∼2%) cCREs were sequence specific only to humans, and 204,921 (∼53%) were with orthologous sequences in the three other species (mammal level 0). An additional 127,301 (∼33%) were orthologous only among three primates (primate level 0) (Fig. 3b, Supplementary Table 8). The number of epi-conserved cCREs dropped with increasing conservation level. Of the 204,921 mammal level 0 cCREs, 66,781 (32.5%) were classified as level 1, 49,135 (24.0%) as level 2, and 16,068 (7.8%) as level 3 epi-conservation (Fig. 3b-d, Extended Data Fig. 4a). Of the 127,301 primate level 0 cCREs (Fig. 3b), we found 60,515 as level 1, 29,395 as level 2, and 32,874 as level 3 epi-conserved in primates but not in mice (Fig. 3b-d, Supplementary Tables 8,9). Finally, 10,743 cCREs showed human-biased accessibility in one or more cortical cell types compared to all three other mammals, and an additional 12,091 human-biased cCREs in primate level 0 sequence regions (Fig. 3d, Supplementary Table 10).

A large proportion of epi-conserved cCREs were found to be promoter-proximal (<1kb from a TSS), and this number increased along with conservation level for both mammal and primate comparisons (Fig. 3e, Supplementary Table 8). Strikingly, mammal level 3 conserved cCREs have a considerably higher fraction of promoter-proximal cCREs than the primate level 3 conserved cCREs. This suggests that the turnover rates are different between the promoter-proximal and distal cCREs during evolution, where distal cCREs evolve much faster than proximal cCREs. Furthermore, the level 3 conserved cCREs that are promoter distal have reduced cell type specificity (Fig. 3e,f), suggesting that distal cCREs with broad cell type activity evolve slower than the cell type-specific distal cCREs.

Transposable elements (TEs) have been proposed as a driver of genomic diversity as they can harbor *cis*-regulatory elements^11, 30^. To characterize the extent of TE contribution to epi-divergence, we calculated the percentage of cCREs within TEs from different conservation levels (Fig. 3g, Supplementary Table 8). We found 75% of human-specific cCREs, and 16% of human-biased cCREs are located within TEs. In contrast,

<1% of mammal level 3 cCREs are located within TEs, highlighting the degree of TE contribution to the species-specific regulatory landscape in human cortical cells. Particularly, L1 and L2 LINEs are the most common TEs containing cCREs, which are most active in excitatory neurons (Fig. 3h). However, human-specific cCREs in different cell types are enriched in different types of TEs. Human-specific cCREs from excitatory neurons (except for L5/6 NP and L6 CT) are most enriched in L1 (Fig. 3h). Glial cells had the strongest enrichment in ERV1 and ERVK LTRs and are lower in L1. While mouse cCREs overlapping TEs show similar trends between conservation groups as humans (Extended Data Fig. 4b), we found enrichment of mouse cCREs in different types of LTRs and SINEs than observed in humans (Extended Data Fig. 4c). Our results provide further evidence that organisms may co-opt TEs to achieve species-specific and cell type-specific gene regulation.

### Comparative analysis of DNA methylomes across species

We further investigate epigenetic evolution by examining differentially methylated regions (DMRs) across all four species. Previous studies have indicated that DMRs are enriched for cCREs^21, 31^. For each species, we called differentially CG methylated regions between cell types using methylpy^32^, identifying 1,361,958 human, 1,661,598 macaque, 1,066,980 marmoset, and 1,748,945 mouse DMRs (Extended Data Fig. 3a,b, Supplementary Tables 11-14). We then identified conserved DMRs across species by repeating the same analysis described for ATAC cCREs.

In terms of conserved sequences, we identified 54,829 human-specific DMR sequences (4.0%), 579,026 (42%) sequences were orthologous in all four mammalian species (mammal level 0), and 519,456 (38%) of sequences found in all three primates but not in mouse (Supplementary Table 15). Of the mammal level 0 DMRs, we find 195,435 (14.3%) as level 1, 144,156 (10.6%) as level 2, and 23,414 (1.72%) as level 3 (Fig. 4a,b). Primate epi-conserved elements were identified from the mammal and primate level 0 cCREs (Fig. 4a). We found an additional 201,415 (14.8%) as level 1, 199,555 (14.65%) as level 2, and 64,138 (4.7%) as level 3 epi-conserved in primates but not in mice, (Fig. 4a,b, Supplementary Tables 15,16).

**Fig. 4:**
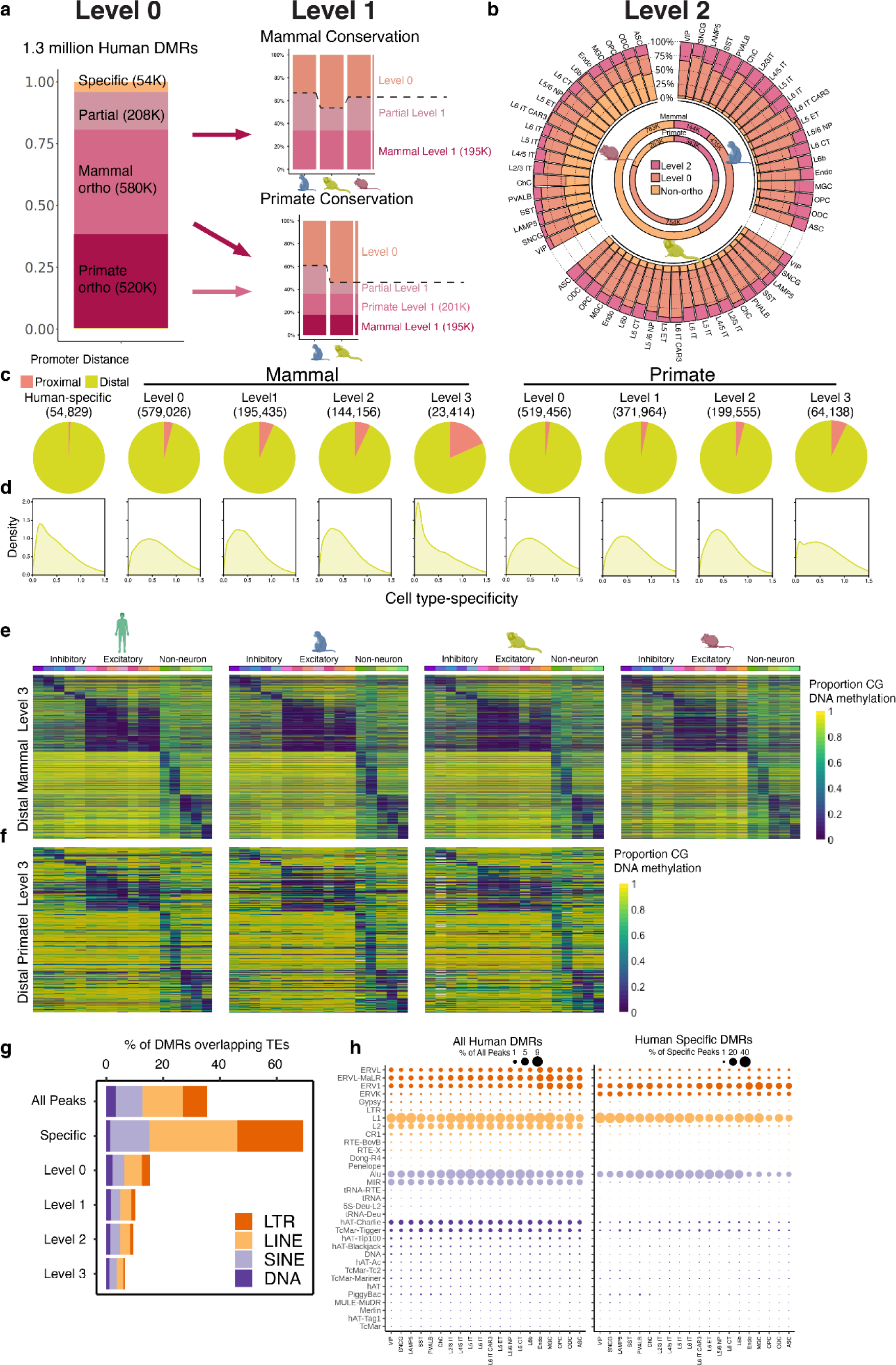
Comparative analysis of DNA methylomes across species. **a.** Proportions of level 0 (sequence conserved) and level 1 (tissue conserved) DMRs across mammals and primates. **b.** Level 2 conservation of human DMRs showing the overlap between each species for the same cell type (outer circle stacked bars). Inner circles show the breakdown for mammal and primate comparisons for all human DMRs. **d.** Kernel density plots showing the cell type specificity of DMRs across cell types. **e.** Heatmaps showing normalized methylation levels among mammal level 3 distal DMRs, ordered by the cell type with the lowest human methylation signals. **f.** Heatmaps showing methylation levels across Primate level 3 DMRs ordered by the cell type with the lowest human methylation signals. **g.** Stacked bar plots showing percentage of DMRs overlapping TEs for different conservation groups. **h.** Dot plots showing the percentage of all (left) or human-specific (right) DMRs overlapping different subclasses of TEs for each cell type.

Compared to cCREs identified using ATAC-seq, DMRs are much less promoter-proximal (Fig. 4c). However, they still displayed an increasing enrichment for TSS proximity with higher levels of conservation. Much like for cCREs identified using ATAC-seq (Fig. 3e), elevated levels of conservation correlate with lower cell-type specificity (Fig. 4d,e,f). Diverging from this trend, human-specific DMR sequences are less cell-type specific (Fig. 4d). Reconciling this difference, we find human-sequence specific DMRs which overlap ATAC-peaks showed increased cell-type specificity compared to those that do not (Extended Data Fig. 5c).

We again evaluated the contribution of TEs to DNA methylation divergence by evaluating the proportion of TEs across different levels of conservation. DMRs showed the same pattern of decreasing TE enrichment with increasing conservation, with human-specific sequences showing enrichment for TEs (69%), and level 3 conserved sequences showing depletion (6.5%) (Fig. 4g, Supplementary Table 15). To evaluate the contribution of TEs to DMRs across cell types we identified the proportion of various TE classes to hypomethylated DMRs in each cell type (Fig. 4h). DMRs show greater enrichment of TE elements than ATAC peaks, with some cell types deriving almost 10% of hypo-DMRs from L1 elements. While both ATAC cCREs and DMRs show enrichment of L1 elements, the enrichment among DMRs shows less cell-type specificity. In contrast, DMR ALU elements show modest enrichment among excitatory neurons which was absent among ATAC cCREs. Human-specific DMRs showed increased enrichment in neuronally hypomethylated ALU elements (Fig. 4h). Like ATAC, human-specific DMRs showed depletion in certain classes of elements, and a notable increase in ERVK elements.

### Comparative analysis of the 3D genome features across species

The genome is organized into topologically associating domains (TADs) that influence gene expression through constraining chromatin contacts between promoters and *cis*-regulatory elements^33^. To characterize the conservation of 3D genome organization, we compared TAD boundary elements across species (Fig. 5a, Supplementary Tables 17-20). Most human Level 0 boundaries were Level 1 conserved in macaque, marmoset, and mouse. 40% (5,118) were tissue conserved in all four species (mammal level 1) (Fig. 5b), and 10% (1,290) conserved in the same cell types (mammal level 2) (Fig. 5c, Supplementary Table 21). Among all cell types, L2/3, L4/5, L5, L6 IT excitatory neurons had the highest number of conserved boundaries across each species (Fig. 5c). 859 human boundaries were sequence-specific to only humans (Fig. 5b) and 1,653 were human-biased. Importantly, we found that genes near conserved boundaries have higher conservation in expression patterns than genes near divergent boundaries (Fig. 5d). cCREs and DMRs within conserved boundaries similarly have more conserved epigenetic states than those within divergent boundaries (Fig. 5d). Our results suggest a co-evolution of the 3D genome along with epigenome and gene expression.

**Fig. 5:**
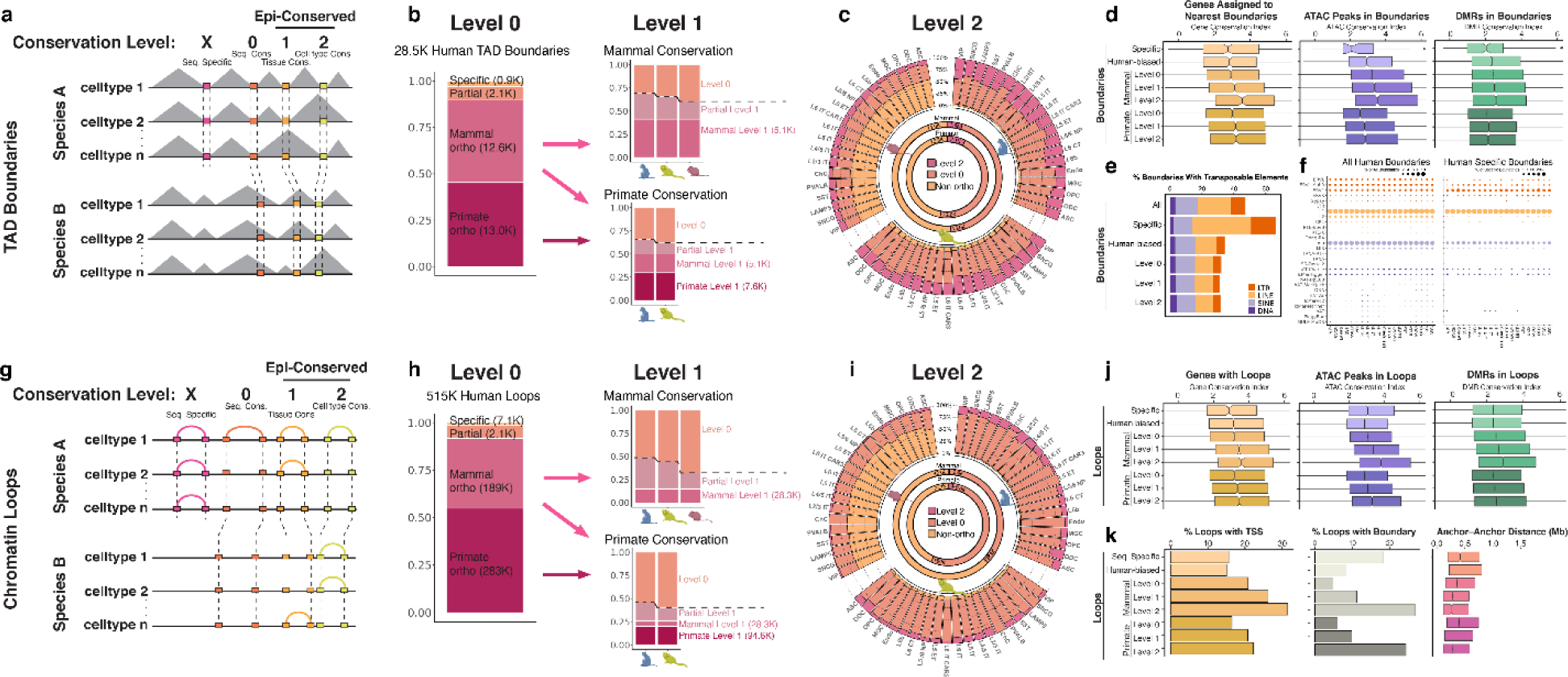
Comparative analysis of TAD boundaries and chromatin loops across species. **a**, Schematic illustrating levels of conservation for TAD boundaries. **b**, Stacked bar plots representing the breakdown of human boundaries for each indicated group for Level 0 and Level 1 conservation. **c**, Level 2 conservation of human boundaries showing the overlap between each species for the same cell type (outer circle stacked bars). Inner circles show the breakdown for mammal and primate comparisons for all human boundaries. **d**, Conservation Index of gene expression, ATAC-seq peaks, and DMRs associated with boundaries of indicated conservation level. **e**, Stacked bar plots showing percentage of boundaries overlapping TEs for different conservation groups. **f**, Dot plots showing the percentage of all (left) or human-specific (right) overlapping different subclasses of TEs for each cell type. **g**, Schematic illustrating levels of conservation for chromatin loops. **h**, Stacked bar plots representing the breakdown of human loops for each indicated group for Level 0 and Level 1 conservation. **i**, Level 2 conservation of human loops showing the overlap between each species for the same cell type (outer circle stacked bars). Inner circles show the breakdown for mammal and primate comparisons for all human boundaries. **j**, Conservation Index of gene expression, ATAC-seq peaks, and DMRs overlapping at least one anchor of loops for each indicated conservation level. **k**, Percent loops with TSS or boundary overlapping at least one anchor bin, or anchor to anchor distance for loops of indicated conservation level.

To determine if TEs are differentially enriched between divergent and conserved boundary elements, we calculated the percentage of boundaries harboring TEs for each conservation group. Strikingly, 67% of human-specific boundary elements contain TEs; by contrast, 32% of mammal level 2 boundaries contain TEs (Fig. 5e). Particularly, L1 and Alu had the highest enrichment in human boundaries, and human-specific boundaries were selectively more enriched for ERV1 TEs (Fig. 5f). Previously, it has been reported that TEs, especially Alu and SINE elements harboring CTCF binding sites may be a mechanism for the evolution of chromatin organization in different species^34^. More recently, several studies showed that the LTR family of TEs also induces the formation of TAD boundaries in specific cell types and developmental stages^35, 36^.

We next classified chromatin loops by conservation levels (Fig. 5g, Supplementary Tables 22-25). Compared to boundary elements, a lower fraction of loops were identified as conserved across mammals and primates (Fig. 5h,i, Supplementary Table 26). Conserved loops were more likely to contain promoters with conserved expression and cCREs and DMRs with conserved activity (Fig. 5j), suggesting that conserved 3D chromatin interactions maintain the conservation of gene regulatory functions. We further characterized loops in each conservation group by calculating the percentage overlap with TSSs and boundaries. Conserved loops had a higher percentage containing TSSs and a higher percentage containing boundaries (Fig. 5k). Additionally, anchor-to-anchor loop distance decreased with increasing conservation group (Fig. 5k), suggesting shorter distance loops are more likely to be preserved through evolution, potentially due to a greater chance that both anchors will be retained in the same syntenic region. Our analysis demonstrates the striking concordance between the conservation of the 3D genome with gene regulatory programs, suggesting selective pressure on genome organization maintains conserved gene regulation throughout mammalian evolution.

### Epigenetic conservation at cCREs is correlated with conservation in expression of their putative target genes

We next ask how epigenetic divergence at cCREs correlates with the evolution of gene expression programs in different species. We first predict the putative target genes for promoter-distal cCREs and likely enhancers with the “Activity-By-Contact” (ABC) model^37^, using our chromatin accessibility and chromatin contact data for each cell type (Fig. 6a, Supplementary Tables 27-30). ABC scores of promoter distal cCREs were highest for mammal level 3 compared with all and human-biased distal cCREs (Fig. 6b), suggesting that conserved cCREs are more likely to function as enhancers. Expression of mammal level 3 distal cCRE target genes is more conserved than human-biased distal cCRE target genes (Fig. 6c), providing evidence that conservation of the enhancer landscape promotes conservation of gene expression during evolution. In contrast, distal human-specific cCREs have a significantly lower percentage of putative enhancers (16.8%) compared to all distal cCREs (25.5%), while the percentage of distal human-biased putative enhancers (34.0%) is significantly higher (Fig. 6d). In general, cCREs located in TEs are less often predicted to function as enhancers (Extended Data Fig. 7). Strikingly, 75.5% of distal mammal level 3 cCREs are predicted to function as enhancers (Fig. 6d). Although human-biased cCREs are less often predicted as enhancers compared to epi-conserved cCREs (Fig. 6d), their putative target genes have higher expression in humans (Fig. 6e), giving evidence for their function as enhancers for human-biased gene expression.

**Fig. 6:**
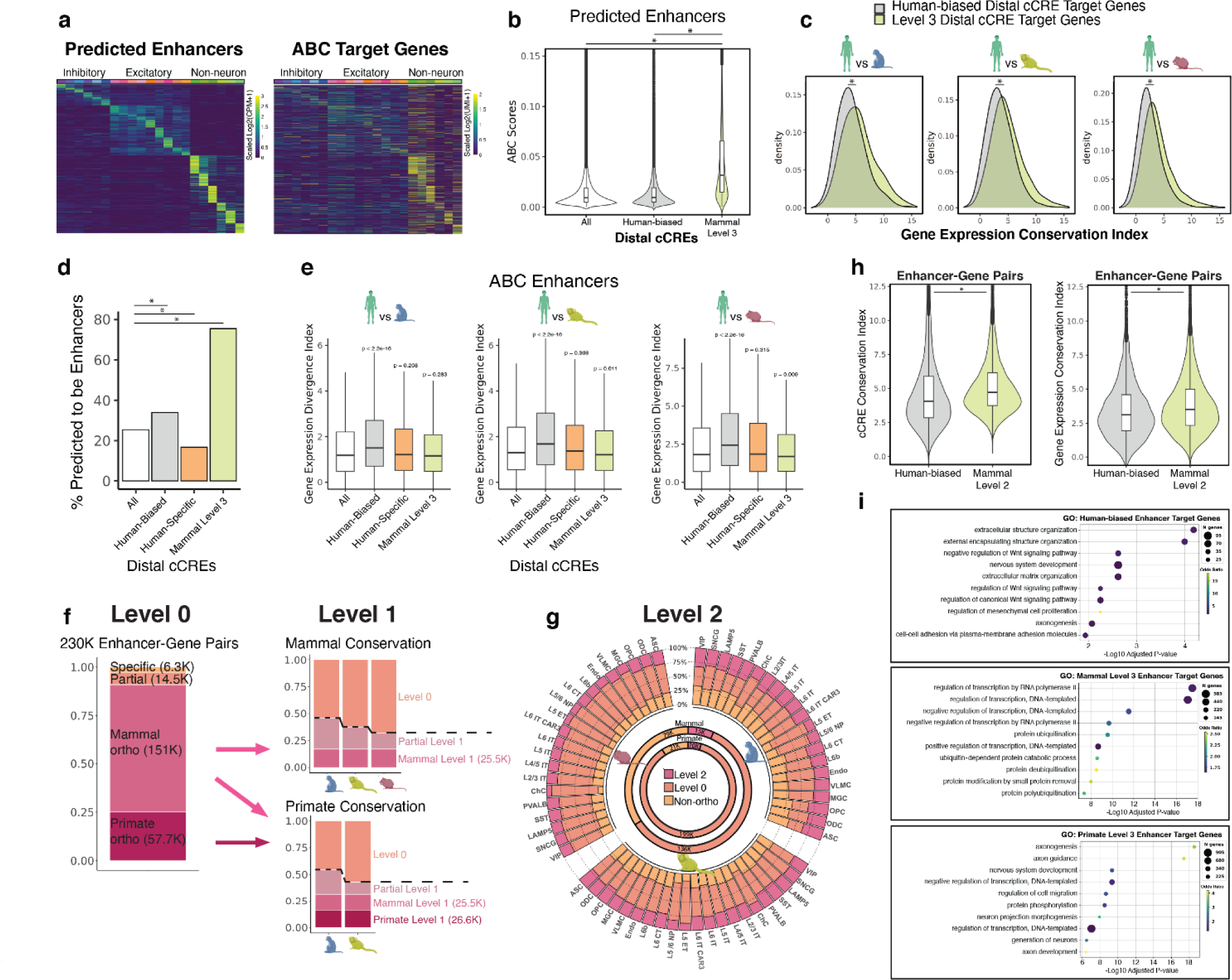
Epigenetic conservation at cCREs is correlated with conservation in expression of their putative target genes. **a**, Heatmaps displaying correlation of activity between predicted enhancers with highest ABC score for each gene with prediction (n=8,083), row scaled. **b**, Violin and box plots of highest ABC score (at least 0.02) across cell types for each distal cCRE from indicated conservation group. * p < 2.2e-16 Wilcoxon rank sum test. **c**, Density plots for gene expression conservation index values from each indicated mammal pairwise comparison with humans. Target genes are separated into two groups: targets of human-biased distal cCREs or mammal level 3 distal cCREs. * p < 2.2e-16 Wilcoxon rank sum test. **d**, Bar plots representing percentage of peaks predicted to be enhancers (ABC score ≥ 0.02). * p < 2.2e-16 Wilcoxon rank sum test. **e**, Box plots of ABC putative target genes for each distal cCRE from indicated conservation group. p-values are from Wilcoxon rank sum test. **f**, Stacked bar plots representing the breakdown of human ABCpredictions for each indicated group for Level 0 and Level 1 conservation. **g**, Level 2 conservation of human ABC predictions showing the overlap between each species for the same cell type (outer circle stacked bars). Inner circles show the breakdown for mammal and primate comparisons for all human boundaries. **h**, Distributions of conservation index values for ATAC-seq peaks (cCREs) or genes involved in ABC enhancer-gene pairs. * p < 2.2e-16 Wilcoxon rank sum test. **i**, Top significant GO analysis Biological Process terms for human-biased enhancer target genes, mammal level 3 enhancer target genes, or primate level 3 enhancer target genes.

**Fig. 7:**
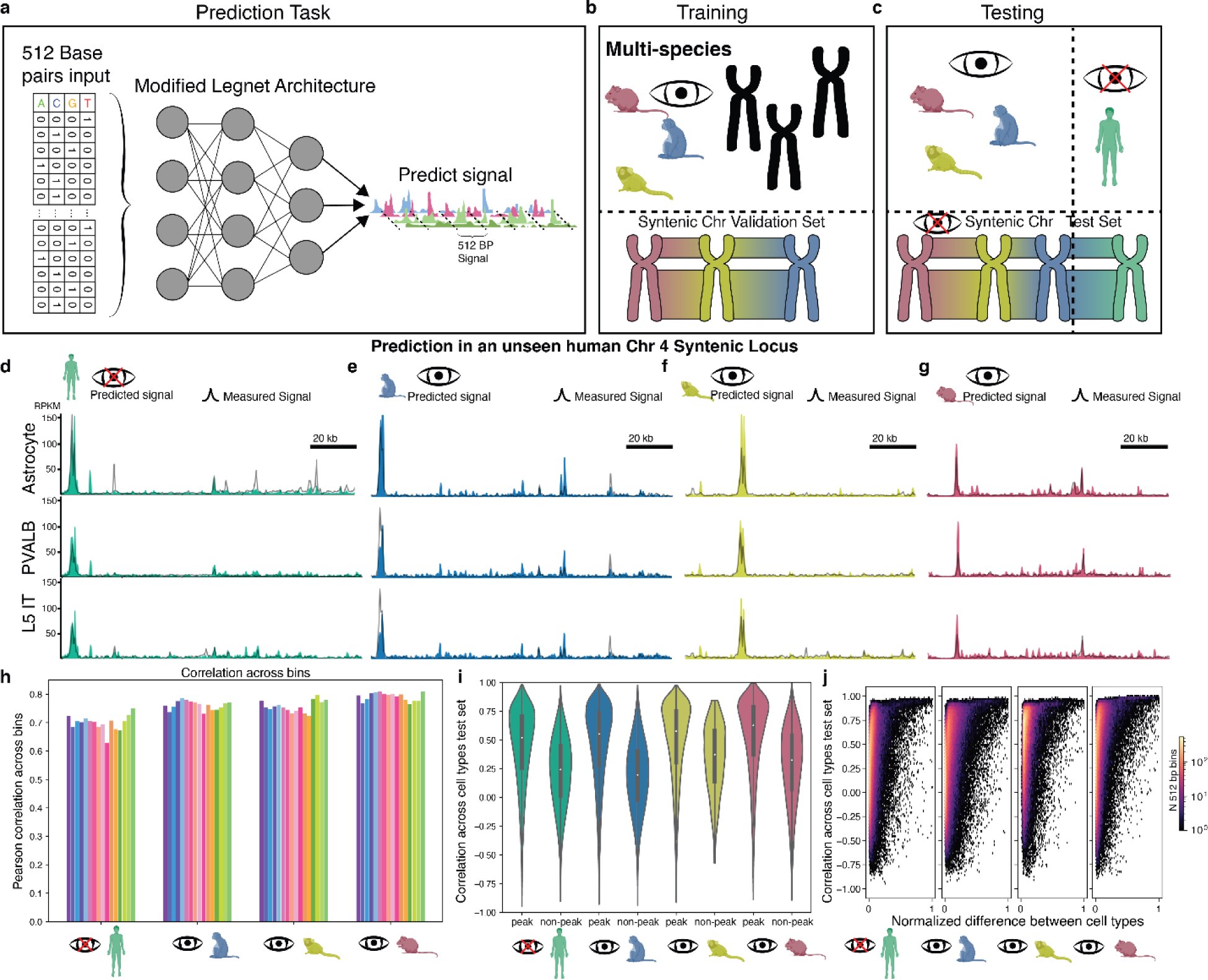
Deep learning models predict cell-type specific chromatin accessibility from DNA sequence alone. **a.** Schematic illustrating the prediction task used to predict cCREs from DNA sequence. **b.** Schematic illustrating the training procedure of the deep neural network. Data from three species are used as training sets, leaving out a set of syntenic chromosomes to be used as testing datasets. **c.** Schematic illustrating the evaluation of the model on an unseen set of syntenic regions in a species excluded from the training dataset. **d.** Model predictions across cell types in the held-out species on a syntenic test set region near the POLARB2 TSS. **e.** Predicted signal across cell types in the same test set region in a species included in Macaque. **f.** Predicted signal across cell types in the same test set region in Marmoset. **g.** Predict chromatin accessibility signal in the same test set region in Mouse. **h.** Correlation of predicted chromatin accessibility to true chromatin accessibility across bins in each species colored by cell type. **i.** Correlation across cell types for each species in both regions with a peak and regions outside of a peak. **j.** Relationship between correlation across cell types and the difference in accessibility across cell types at a given locus.

We classified ABC-predicted human enhancer-gene pairs into conservation levels and identified 25,472 mammal level 1 pairs and 15,142 mammal level 2 pairs (Fig. 6f,g, Supplementary Table 31). In general, cCREs and genes in mammal level 2 enhancer-gene pairs were more conserved than those in human-biased enhancer-gene pairs (Fig. 6h). Consistent with our gene expression conservation analysis, we found that target genes of mammal level 3 cCREs tend to be enriched for those involved in transcriptional regulation, and target genes of primate level 3 cCREs are enriched for nervous system and neuronal functions (Fig. 6i). Our comparative analysis of enhancer-gene pairs in M1 cells indicates the extent of conservation of the enhancer regulatory landscape across mammals and primates.

### Deep learning models predict cell-type specific chromatin accessibility from DNA sequence alone

Previous studies have suggested a conserved regulatory grammar and syntax at cis regulatory elements in the genomes of mammalian species^38^, but how the genome encodes the gene regulatory program continues to be elusive. To understand how differences in species chromatin accessibility are driven by sequence changes, we trained a neural network model to predict the chromatin accessibility of a short region (512 base pair or bp) in each cell type from the DNA sequence alone (Fig. 7a, Extended Data Fig. 8a). We adapt Legnet^39^ to this task which has achieved state of the art prediction accuracy for short sequence MPRA activity. We trained our model on three species and evaluated on a fourth unseen species (Fig. 7b,c). To avoid data leakage ^40^, we evaluated the performance of our model on a held-out test set of largely syntenic chromosomes, mitigating the risk that an orthologous region had been seen in the dataset on another species (Fig. 7c).

Our model is able to predict chromatin accessibility in our testing dataset regardless of the species inclusion in the training data (Fig. 7h), consistent with the notion of a general conservation of grammar and syntax of regulatory elements. This is in line with previous observations on the tissue level^41, 42^. Our model also demonstrates cell type specificity in its predictions on the test set showing a 0.533 median correlation across cell types in peak regions for the held-out human dataset (Fig. 7i). The cell type discriminatory ability of our model generally increased with the difference between the most accessible and least accessible cell type for a given region (Fig. 7j). The results demonstrate a conserved gene regulatory grammar across mammalian neocortical cell types, and showcase the considerable ability to identify cis-regulatory sequences with a small receptive field.

### Leveraging epigenome conservation to interpret disease risk genetic variants

Genome-wide association studies (GWAS) have identified common genetic variants linked to various traits and diseases, yet most GWAS variants reside in non-coding regions of the genome and their influence on gene expression remains unresolved^43^. A growing body of evidence suggests that the noncoding disease risk variants may contribute to disease by disrupting *cis* regulatory elements and affecting gene expression in cell types relevant to disease pathogenesis^44–46^. Since human cCREs with elevated epigenetic conservation levels are more likely to be predicted as active enhancers, we hypothesize that evidence of epigenetic conservation may improve our ability to interpret noncoding disease risk variants. To test this, we performed stratified linkage disequilibrium score regression (S-LDSC) analysis^47^. When this analysis was performed with all the cCREs, we observed high enrichment for variants implicated in neurological traits within cCREs identified from various neuronal and glial cell types, as expected (Fig. 8a). However, when this analysis was performed with the human epi-divergent cCREs (specific and human-biased cCREs), the enrichment for GWAS variants associated with neurological traits is virtually eliminated (Fig. 8a). By contrast, the enrichment improves when the analysis is carried out with the mammal level 2 epi-conserved cCREs (Fig. 8a,b). For example, multiple sclerosis (MS) associated variants are highly enriched for epi-conserved microglia cCREs, but not significantly enriched in any cell type when considering the full set of cCREs (Fig. 8a,c). Two other examples include anorexia nervosa and tobacco use disorder, which only show significant enrichments in neuronal epi-conserved cCREs (Fig. 8a,c).

**Fig. 8:**
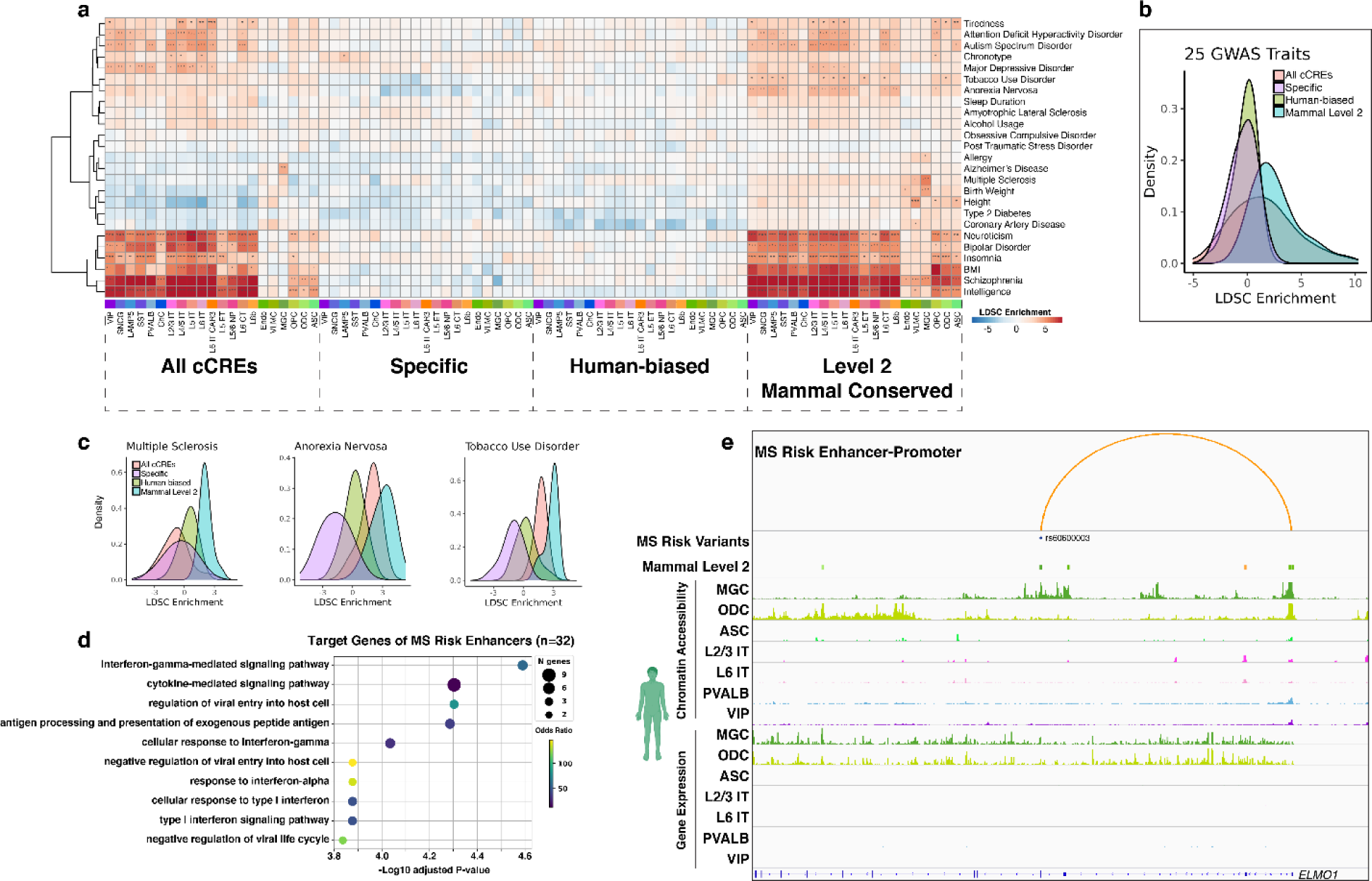
Leveraging epigenetic conservation for interpreting non-coding risk variants of neurological disease and traits. **a**, Linkage disequilibrium score regression analysis to identify GWAS enrichments in cCREs of each cell type for different conservation sets. * FDR < 0.001, ** FDR < 0.0001, *** FDR < 0.00001. **b**, Distribution of LDSC enrichments across cells for each of the 25 traits from **a**. **c**, Distribution of LDSC enrichments across cells for multiple sclerosis, anorexia nervosa, and tobacco use disorder. **d**, Top significant GO analysis Biological Process terms for ABC target genes of enhancers harboring a multiple sclerosis risk variant^48^. **e**, Example locus of a mammal level 2 predicted enhancer of *ELMO1* overlapping a multiple sclerosis risk variant in a microglia-specific chromatin accessible region.

Since our list of epi-conserved cCREs specifically linked microglia regulatory elements to MS, we used our enhancer-gene predictions to see if we could interpret potential gene regulatory effects of MS risk variants and their relation to microglia functions. Using a list of 233 MS risk variants^48^, we identified 38 overlapping human cCREs with 32 predicted target genes. The target genes of enhancers harboring MS risk variants are enriched for functions related to immune response pathways (Fig. 8d). For example, MS risk variant rs60600003 resides in a mammal level 2 cCRE in the intron of *ELMO1*, a gene involved in phagocytosis (Fig. 8e). *ELMO1* is expressed in both microglia and oligodendrocytes; however, the cCRE containing the risk variant rs60600003 is accessible exclusively in microglia (Fig. 8e). This microglia specific cCRE is predicted to be an enhancer of *ELMO1*, implicating *ELMO1* expression to be selectively affected in microglia by this MS risk variant. Our analysis provides examples of how comparative epigenomics can help with interpreting disease-risk genetic variants for neurological diseases such as MS.

## Discussion

Our comparative analysis of the transcriptome, epigenome, and 3D genome features of 21 cortical cell types from four species provides an unprecedented perspective on the evolution of gene regulatory programs in rodents and primates. Through integrating four molecular modalities (gene expression, chromatin accessibility, DNA methylation and chromatin conformation) across 21 cell types in four species, we characterized conserved and divergent gene regulatory features focusing on three evolutionary times scales: mammal conserved (from human to mouse, ∼90M years), primate conserved (from human to marmoset, ∼43M years), and human divergent (human from macaque, ∼25M years ago). Although their turnover rates are slower than neutrally evolving sequences, we find that the epigenetic state at the majority of cCREs reported in the present study is not conserved in mammalian evolution and their selective constraint is dependent on the type of *cis*-regulatory element. For example, epigenetic conservation of distal cCREs is generally lower than that of promoters or promoter-proximal cCREs, and epigenetic conservation of cCREs with high cell type specificity is lower than those with broad cell type activities, consistent with previous findings^4–7^. Compared to epi-divergent distal cCREs, we show that epi-conserved distal cCREs are more often predicted to act as enhancers of target genes (Fig. 6b) and are more enriched for genetic variants associated with neurological disease/traits (Fig. 8a). Taken together, this provides evidence that comparative epigenomics can assist in identifying functional enhancers. In addition, our data provide evidence that selective pressure on 3D genome organization maintains conserved gene regulatory programs (Fig. 5d,j).

We demonstrate new evidence that TEs may be a major source of species-specific cCREs (Fig. 3g), DMRs (Fig. 4g), and TAD boundaries (Fig. 5e). Notably, different types of TEs contribute to the establishment of different categories of cCREs. For example, ERVK contributes highly to human-specific cCREs (Fig. 3h, Extended Data Fig. 7a,b) but not to human-specific boundaries (Fig. 5f). TE contribution is also cell type dependent. cCREs in L1 are more active in excitatory neurons while human-specific cCREs in ERV1 and ERVK have higher accessibility in glia (Fig. 3h). This observation suggests a role for TEs in the regulation of cell type-specific gene expression. We highlight the power of machine learning approaches to learn the gene regulatory grammar from single-cell multi-omic datasets across mammalian species and multiple cell types (Fig. 7, Extended Data Fig. 8c,d). The sequence-based predictors resulting from the model training demonstrate a remarkable predictive ability, suggesting a general conservation in regulatory grammar across mammalian species. Our results also support the notion that epigenetic divergence is primarily driven by sequence divergence. While neural networks have shown promise in predicting epigenetic features and gene expression levels from DNA sequence^39, 41, 49^, there is still a gap between current approaches and experiment-level predictions. While recent advances have been considerable, work in neural network scaling suggests improvements in model accuracy grow following a power law, requiring an exponential increase in both model and dataset size^50^. This need for increasing data presents a significant hurdle in achieving highly accurate sequence-based prediction of gene expression. The human genome alone provides a limited sequence set to overcome these challenges, but they can be met by expanding the repertoire of species in epigenomic datasets.

## Methods

### Nuclei preparation from frozen brain tissue for Chromium Single Cell Multiome ATAC + Gene Expression (10x Genomics)

M1 tissue was obtained from three human donors (42 y.o. male, 29 y.o. male, and 58 y.o. male), three macaque donors (6 y.o. male Macaca mulatta, 6 y.o. male Macaca mulatta, and 14 y.o. male Macaca fascicularis), three marmoset (Callithrix jacchus) donors (5 y.o. male, 4 y.o. male, and 6 y.o. female), and MOp from eight P56 C57BL/6J male mice (Mus musculus). Mouse MOp was dissected into four subregions (2C, 3C, 4B, 5D) as described in Li et al. 2021^19^. Each subregion was pooled from four mice for each replicate, and a total of two replicates was performed for each subregion.

Brain tissue was pulverized using a mortar and pestle on dry ice and pre-chilled with liquid nitrogen. Pulverized brain tissue was resuspended in 1mL of chilled NIM-DP-L buffer (0.25M sucrose, 25mM KCl, 5mM MgCl2, 10mM Tris-HCl pH 7.5, 1mM DTT, 1X Protease Inhibitor (Pierce), 1U/μL Recombinant RNase inhibitor (Promega, PAN2515), and 0.1% Triton X-100). Tissue was Dounce homogenized with a loose pestle (5-10 strokes) followed by a tight pestle (15-25 strokes) or until the solution was uniform.

Nuclei were filtered using a 30μm CellTrics filter (Sysmex, 04-0042-2316) into a LoBind tube (Eppendorf, 22431021) and pelleted (1000 rcf, 10 min at 4°C) (Eppendorf, 5920 R). Pellet was resuspended in 1mL NIM-DP buffer (0.25M sucrose, 25mM KCl, 5mM MgCl2, 10mM Tris-HCl pH 7.5, 1mM DTT, 1X Protease Inhibitor, 1U/μL Recombinant RNase inhibitor) and pelleted (1000 rcf, 10 min at 4°C). Pelleted nuclei were resuspended in 400uL 2μM 7-AAD (Invitrogen, A1310) in Sort Buffer (1mM EDTA, 1U/μL Recombinant RNase inhibitor, 1X Protease Inhibitor, 1% fatty acid-free BSA in PBS). 120,000 nuclei were sorted (Sony, SH800S) into a LoBind tube containing Collection Buffer (5U/uL Recombinant RNase inhibitor, 1X Protease Inhibitor, 5% fatty acid-free BSA in PBS). 5X Permeabilization Buffer (50mM Tris-HCl pH 7.4, 50mM NaCl, 15mM MgCl2, 0.05% Tween-20, 0.05% IGEPAL, 0.005% Digitonin, 5% fatty acid-free BSA in PBS, 5mM DTT, 1U/μL Recombinant RNase inhibitor, 5X Protease Inhibitor) was added for a final concentration of 1X. Nuclei were incubated on ice for 1 minute, then centrifuged (500 rcf, 5min at 4C). Supernatant was discarded and 650uL of Wash Buffer (10mM Tris-HCl pH 7.4, 10mM NaCl, 3mM MgCl2, 0.1%.Tween-20, 1% fatty acid-free BSA in PBS, 1mM DTT, 1U/μL Recombinant RNase inhibitor, 1X Protease Inhibitor) was added without disturbing the pellet followed by centrifuging (500 rcf, 5 min at 4°C). Supernatant was removed, and the pellet was resuspended in 7uL of 1X Nuclei Buffer (Nuclei Buffer (10x Genomics), 1mM DTT, 1 U/μL Recombinant RNase inhibitor). 1μL of nuclei was diluted in 1X Nuclei Buffer, stained with Trypan Blue (Invitrogen, T10282) and counted. 16-20k nuclei were used for tagmentation reaction and controller loading and libraries were generated following manufacturer’s recommended protocol (https://www.10xgenomics.com/support/single-cell-multiome-atac-plus-gene-expression). 10x multiome ATAC-seq and RNA-seq libraries were paired-end sequenced on NextSeq 500 and NovaSeq 6000 to a depth of ∼50,000 reads per cell for each modality.

### Genome Assemblies and Annotations

*Homo sapiens (Human)* assembly: hg38 GRCh38 annotation: hg38 Gencode v33

*Mus musculus (Mouse)* assembly: mm10 GRCm38 annotation: mm10 Gencode vM22

*Macaca mulatta (Rhesus monkey)*

assembly: Mmul_10 (rheMac10)

annotation: ensembl release 104 (and Refseq GCF_003339765.1 for 10x multiome (see below))

*Callithrix jacchus (white-tufted-ear marmoset)*

assembly: cj1700_1.1 (calJac4) annotation: GCA_009663435.2

To maximize the number of orthologous protein-coding quantified in macaque 10x multiome RNA data, we supplemented any missing protein coding genes in GCF_003339765.1 gtf with annotations present in Ensembl release 104.

### 10x multiome sequence data processing and clustering

Raw sequencing was processed using cellranger-arc (10x Genomics), generating snRNA-seq UMI count matrices for intronic and exonic reads mapping in the sense direction of a gene. We performed unsupervised clustering with RNA UMI counts using Seurat v4^53^ standard analysis pipeline. First cells were filtered for low quality nuclei by requiring ≥1000 ATAC fragments and ≥500 genes detected per nuclei. Counts were normalized using SCTransform identifying 3,000 variable genes used for principal component analysis (PCA). Putative multiplets were predicted using DoubletFinder^54^ software and 10% of cells were removed from each sample that had the highest doublet score. Batch correction across donors was performed using Harmony^55^ on SCTransformed PCs. A k-nearest neighbors graph was built using PCs 1:20 and clusters were identified using Louvain clustering. To visualize clusters, we performed the non-linear dimension reduction technique, UMAP^52^. We annotated subclass-level cell types for mouse, marmoset, and human cells by reference mapping to published M1 snRNA-seq datasets^17, 18^ using Seurat. We integrated datasets from all four species using reciprocal PCA using genes orthologous in all four species as integration anchors, which projects each species datasets into the others’ PCA space and identifies anchors by the same mutual neighborhood requirement. For integration anchors, we only considered genes that are orthologous across all four species. Reads from 21 annotated cell types were combined to generate pseudo-bulk datasets used for downstream analyses.

### ATAC-seq peak calling and filtering

We utilized MACS2 for ATAC-seq peak calling on pseudo bulk ATAC-seq fragments using command macs2 callpeak with parameters --shift −75 --ext 150 --bdg -q 0.1 -B --SPMR --call-summits -f BAMPE. We extended the peak summit by 249 bp upstream and 250 bp downstream to achieve 500 bp width for every peak. Since the number of peaks called in each cell type is related to sequence depth which is highly variable due to differences in cell type abundance, we converted MACS2 peak scores (−log10(q-value)) to ‘score-per-million’^56^. Peaks with a score-per-million of ≥ 2 were retained for each cell type. We further filtered human and mouse peaks by removing those with ENCODE blacklist regions https://mitra.stanford.edu/kundaje/akundaje/release/blacklists/ of hg38 and mm10. For comparative analysis of human ATAC-seq peaks, we first removed peaks that were mapping to a region in any of the four species that had low read mappability. To identify regions with low mappability in our ATAC-seq data, we counted all reads in 1 kb bins across each genome. We took 1 kb bins with 0 reads, and for the remaining bins we took the 0.02 quantile for the number of reads mapped and extended by 1 kb in both directions giving us 3 kb low mappability bins. Finally, low mappability bins within 5 kb were stitched together, providing our final list of low ATAC-seq mappability regions. Peaks or orthologous elements falling in any of these regions in any species were excluded for comparative analysis.

### Nuclei isolation and Fluorescence Activated Nuclei Sorting (FANS)

For all snm3C-seq samples, in-situ 3C treatment was done during the nuclei preparation that allows capturing the chromatin conformation modality as described previously^22^. These steps were carried out using the Arima-3C BETA Kit (Arima Genomics). The nuclei were isolated and sorted into 384-well plates using previous methods^21^. Briefly, single-nuclei were stained with AlexaFluor488-conjugated anti-NeuN antibody (MAB377X, Millipore) and Hoechst 33342 (62249, ThermoFisher) followed by FANS using a BD Influx sorter with single-cell (1 drop single) mode.

### Library preparation and Illumina sequencing

The snm3C-seq samples followed the library preparation protocol detailed previously ^21, 22^. This protocol has been automated using the Beckman Biomek i7 instrument to facilitate large-scale applications. The snm3C-seq libraries were sequenced on an Illumina NovaSeq 6000 instrument, utilizing one S4 flow cell per 16 384-well plates and employing a 150 bp paired-end mode.

### Data preprocessing

#### Mapping and quality control of snm3C

The snm3C-seq mapping is conducted using the YAP pipeline (cemba-data v1.6.8), as previously described^21^. Specifically, the main mapping steps include (1) demultiplexing FASTQ files into single cells (cutadapt, v2.10); (2) reads level quality control (QC); (3) mapping (one-pass mapping for snmC, two-pass mapping for snm3C) (bismark v0.20, bowtie2 v2.3); (4) BAM file processing and QC (samtools v1.9, picard v3.0.0); and (5) methylome profile generation (allcools v1.0.8); (6) chromatin contact calling. All reads from human, macaque, marmoset and mouse were mapped to the hg38, Mmul_10, calJac4 and mm10, respectively.

Pre-analysis quality control for DNA methylome cells was (1) overall mCCC level < 0.05; (2) overall mCH level < 0.2; (3) overall mCG level < 0.5; (4) total final reads > 500,000 and < 10,000,000; and (5) Bismarck mapping rate > 0.5. Note the mCCC level serves as an estimation of the upper bound of the cell-level bisulfite non-conversion rate. Additionally, we calculated lambda DNA spike-in methylation levels to estimate the non-conversion rate for each sample. To prevent any meaningful cell or cluster loss, we choose loose cutoffs for the pre-analysis filtering. The potential doublets and low-quality cells were accessed in the clustering-based quality control described below. For the 3C modality in snm3C-seq cells, we also required cis-long-range contacts (two anchors > 2500 bp apart) > 50,000.

#### Methylome Clustering Analysis

After mapping, single-cell DNA methylome profiles of the snm3C-seq datasets are stored in the “All Cytosine” (ALLC) format, which is a tab-separated table compressed and indexed by bgzip/tabix. The “generate-dataset” command in the ALLCools package can help generate a methylome cell-by-feature tensor dataset (MCDS), stored in Zarr format. We used non-overlapping chromosome 100-kb (chrom100k) bins of the corresponding reference genome to perform clustering analysis, gene body regions ±2 kb for clustering annotation and integration with the companion 10x multiome dataset. Details about the integration analysis are described in the following section.

#### Methylome Clustering

We then perform clustering on the chrom100k matrices. The clustering analysis within each iteration is described in the previous study. In summary, the clustering process includes the following main steps:

1. Basic feature filtering based on coverage and ENCODE blacklist.
2. Highly Variable Feature (HVF) selection.
3. Generation of posterior chrom100k mCH and mCG fraction matrices, as used in the previous study and initially introduced by Smallwood et al.^57^
4. Clustering with HVF and calculating Cluster Enriched Features (CEF) of the HVF-clusters. This framework is adapted from cytograph2 developed by La Manno et al.^58^. We first perform clustering based on variable features and then use these clusters to select CEFs with stronger marker gene signatures of potential clusters. The concept of CEF is introduced in Zeisel et al^59^. The calculation and permutation-based statistical tests for calling CEFs are implemented in “ALLCools.clustering.cluster_enriched_features”, where we select for hypo-methylated genes (corresponding to highly-expressed genes) in methylome clustering.
5. Calculation of principal components (PC) in the selected cell-by-CEF matrices and generation of the T-SNE and UMAP^60^ embedding for visualization. T-SNE is performed using the openTSNE^61^ package with procedures described in Kobak and Berens 2019^62^.

### Cluster-level DNA Methylome Analysis

After clustering analysis, we merged the single-cell ALLC files into pseudo-bulk level using the “allcools merge-allc” command. Next, we performed DMR calling as previously described^63^ using methylpy. In brief, we first calculated CpG differential methylated sites (DMS) using a permutation-based root mean square test^63^. The base calls of each pair of CpG sites were added before analysis. We then merged the DMS into DMR if they are (1) within 500 bp and (2) the minimum methylation difference was greater than or equal to 0.3 across samples. We applied the DMR calling framework across the cell clusters in each species.

### Cell and Cluster-level 3D Genome Analysis

#### Generate chromatin contact matrix and imputation

After snm3C-seq mapping, we used the cis-long range contacts (contact anchors distance > 2,500 bp) and trans contacts to generate single-cell raw chromatin contact matrices at three genome resolutions: chromosome 100-Kb resolution for the chromatin compartment analysis; 25-Kb bin resolution is for the chromatin domain boundary analysis; 10-Kb resolution for chromatin loop or dot analysis. The raw cell-level contact matrices are stored in HDF5-based scool format. We then used the scHiCluster package (v1.3.2) to perform contact matrix imputation. In brief, the scHiCluster imputes the sparse single-cell matrix in two steps: the first step is Gaussian convolution (pad=1); the second step is to apply a random walk with restart algorithm on the convoluted matrix. The imputation is performed on each cis-matrix (intrachromosomal matrix) of each cell. For 100-Kb matrices, the whole chromosome is imputed; for 25-Kb matrices, we imputed contacts within 10.05Mb; for 10-Kb matrices, we imputed contacts with 5.05Mb. The imputed matrices for each cell were stored in cool format. For most following analyses, cell matrices were aggregated into cell groups identified in the previous section. These pseudo-bulk matrices are concatenated into a tensor called CoolDS, stored in Zarr format.

### Compartment analysis

We used the imputed cluster-level contact matrices at 100-Kb resolution to perform the compartment analysis. We used cooltools (version 0.5.1) to compute the PC1 values and used the corresponding genome sequences of the four species to adjust the A/B compartment.

### Domain boundary analysis

We used the imputed cell-level contact matrices at 10-Kb resolution to identify the domain boundaries within each cell using the TopDom algorithm^64^. We first filter out the boundaries that overlap with ENCODE blacklist v2.

We used cooltools (version 0.5.1) to call cluster-level boundaries and domains with 10 Kb resolution matrices. A sliding window of 500 Kb was used to compute the insulation score of each bin, and the bins with boundary strength > 0.1 were selected as domain boundaries.

### Loop analysis

We called the cluster-level loops with 10 Kb resolution matrices using the call_loop function in the scHiCluster package.

### Identification of orthologous sequence elements across species (level 0)

We identified orthologous sequences for each human *cis*-regulatory region in all other species, using liftOver^29^. For each human ATAC-peak and DMR, we first performed liftover to each other species’ genome with a requirement of 50% retained sequence identity (minMatch=0.5). For loop anchors and boundaries, lifted-over required only 30% of retained sequence identity (minMatch=0.3) to account for the difficulty of lifting over a longer (10kb) region. Any region that could not be lifted to any of the other profiled species was identified as human-specific. For ATAC peaks (500 bp), we only kept orthologous elements that are 1 kb or less to the lifted-over genome. Afterward, we performed liftOver from the identified orthologous sequence back to the human sequence. We kept all sequences that mapped back to the same peak identity as “level 0 conserved” between humans and the respective species. We then further identified sequences that are level 0 across all mammals and level 0 across primates.

### Identification of human level 1 (tissue conserved) and level 2 (cell type conserved) cis-regulatory elements

For each human feature (DMR, ATAC peak, loop, boundary, and ABC enhancer-pair) we determined if the feature was also present across species. For each non-human species, we used each feature’s orthologous coordinates in hg38 and performed bedtools^65^ intersect^65^, counting each human element with an overlapping element as having level 1 conservation between human and that species independent of cell type. We further identified elements that are level 1 across all mammals (mammal level 1), as well as elements that are level 1 across primates and not in mouse (primate level 1). Elements were identified as level 2 conserved if the intersection, as described above, existed in any of the same cell types across species. This procedure was modified for DMRs, with DMRs being split into hypo-and hyper-DMRs and performing the described procedure for each; the results of both were aggregated. For loops, this procedure was modified by requiring intersection at both anchor bins. For ABC enhancer pairs, we required that the orthologous cCRE targeted the orthologous gene across species.

### Identification of level 3 conserved Peaks and DMRs

For each species pair, we identified ATAC Peaks and DMRs with conserved patterns of activity across cell types. We first normalized peak accessibility in each cluster to Log2 counts per million counts quantified for level 0 mammal peaks or the combined set of mammal and primate level 0 peaks (when comparing primates). For DMRs, we transformed quantifications to 1 - the CG methylation level in each cell type. We then consider the effect size (T-statistic) of a GLS (generalized least squares) regression^25^ between the species as the effect size of conservation. This procedure controls for the effects of dependence between cell types. A key step in generalized least squares is to estimate the covariance matrix. For each species pair, we compute a covariance matrix between cell types by first taking the covariance between cell types for each species across all peaks or DMRs. We then form a covariance matrix for the regression by taking the mean of both species’ covariance. Given the GLS t-statistic for each species pair, we then identify conserved genes between each species with a false discovery rate of 0.05 adjusted using the Benjamini-Yekulti^66^ to account for dependency among cCREs.

We further identify two categories for peaks and DMRs: Those conserved among mammals which were identified as conserved between each pair of species, and those conserved among primates which were identified as conserved among all three primates but not among all species.

### Cell-type specificity of genes, ATAC peaks, and DMRs

For each gene, ATAC peak, and DMR, we computed its cell-type specificity using an information-theoretic criteria^67^. We identified ubiquitously expressed genes as those with a specificity of less than 0.01. For DMRs, we transformed quantifications to 1 - the methylation level in each cell type.

### Annotation of transposable elements and TSS proximity

For each human element in each category (DMR, Peak, Loop, boundary), we annotated its transposable element association and identified its TSS proximity using annotatePeaks.pl with hg38 from HOMER^68^. This analysis was repeated for mouse ATAC peaks using mm10 to identify their transposable element association.

### Gene ontology (GO) enrichment analysis

We performed gene ontology enrichment analysis using the Enrichr^69^ module in GSEApy^70^ The background gene set was all expressed genes. For ABC target genes, the background set was all human genes called as having an ABC enhancer.

### Identification of species-biased gene activity

Starting with a list of one-to-one orthologous genes across all four species, we performed differential expression analysis on pseudo bulk count profiles for each cell type using EdgeR^26^. We performed analysis using the recommendations of ^71^. Each pseudo bulk profile was normalized for sequencing depth using trimmed mean of M-values normalization^72^, after which tagwise dispersion was estimated using “locfit”. We fit a single model to predict the expression of a cell type based on species identity using “glmFit”, after which differential expression was evaluated on between species contrasts for each species pair. We used stringent criteria to identify if a gene is differentially expressed between a species pair. To account for multiple comparisons we nominated a false discovery rate of 0.001, which we further lowered to 8.33 * 10^−6^, by dividing by the number of pairs of species (6), multiplied by the number of cell types (20). In addition to this FDR threshold, we required our DE genes to meet a minimum fold change of 2, as well as be expressed in at least 15% of the cells in the upregulated species cell type.

Following these criteria, we further identified biased genes for each species. For each cell type in each species, we identified biased genes if a gene was significantly upregulated in that cell type when compared to each other species.

### Identification of peaks with species-biased chromatin accessibility

Starting from the sets of human peaks with orthologs in all four species, we used EdgeR to identify differential chromatin accessibility across species. We used the same parameters as used for identifying species-biased gene activity to estimate fold changes and p-values for each orthologous peak region. When identifying significantly differentially accessible peaks, we make some modifications. We use the same FDR cutoff (8.33 * 10^−6^); however, to account for the sparseness of peaks, we no longer place a threshold on the number of cells where a peak was detected. To compensate, we require a minimum fold change between species of at least 4.

Following these significance criteria, we further identified biased peaks for each cell type in each species. For each cell type in each species, we identified a biased peak if the peak was significantly upregulated in that cell type when compared to each other species.

### Identification of genes with conserved patterns of activity

For each species pair we identified genes with conserved patterns of activity across cell types. We first normalized gene expression in each cluster to Log2 counts per million counts quantified in orthologous genes. We then consider the effect size (T-statistic) of a GLS (generalized least squares) regression ^25^ between the species as the effect size of conservation. This procedure controls for the effects of dependence between genes. A key step in generalized least squares is to estimate the covariance matrix. For each species pair, we compute a covariance matrix between cell types by first, taking the covariance between cell types for each species across all genes. We then form a covariance matrix for the regression by taking the mean of both species’ covariance. Given the GLS T statistic for each species pair, we then identify conserved genes between each species with a false discovery rate of 0.05 adjusted using the Benjamini-Yekulti^66^ to account for dependencies among genes.

We further identify two categories for genes: Those conserved among mammals which were identified as conserved between each pair of species, and those conserved among primates which were identified as conserved among all three primates but not among all species.

### Cross-species open chromatin Legnet

We trained a deep learning model to predict open chromatin based on the architecture of Legnet^39^. In short, our model takes as input a 512 base-pair bin of DNA sequence and predicts the log2(RPKM+1) normalized chromatin of accessibility within that 512 base-pair bin, as well as binary peak calls across all cell types.

#### Dataset design

For each species, we generated a dataset of sequences using the union peak set identified across cell types. Additionally, we selected a random control region for each peak in the same chromosome. From this dataset, we selected test and validation datasets through identifying chromosomes with high synteny across species to minimize data leakage^40^. Chromosomes were selected by visualizing region correspondence in the NIH National Library of Medicine Comparative Genome Viewer^73^.

#### Model design

We built legnet using a convolutional kernel size of seven and efficient-net blocks of size [256, 128, 64, 64]. At the final layer, we performed average pooling to a tensor of size 80, followed by four separate linear heads of length 20, one for each species in the training dataset and one which was trained to predict values for all species.

#### Training and loss

While training, we used two losses, first, a Possion Non-Negative-log-likelihood loss between the predicted signal, and the true signal^49^. We also computed MSE loss on the predicted pairwise difference in predicted signal between each cell type and the difference in true signal for each cell type. For each species in the training set, we evaluated the prediction loss on the output head specific to that species, as well as the all species output head. We trained the model for 25 epochs using the AdamW optimizer with a learning rate of 0.002, saving the model with the lowest loss on the validation set.

#### Model evaluation

The model was evaluated on the test chromosome set of each species, as well as the test chromosome set of the unseen species.

### External datasets

PhastCons^74^ conserved elements were downloaded from the UCSC genome browser (http://hgdownload.cse.ucsc.edu/goldenpath/mm10/phastCons60way/).

## Statistics

No statistical methods were used to predetermine sample sizes. There was no randomization of the samples, and investigators were not blinded to the specimens being investigated. Low-quality nuclei and potential barcode collisions were excluded from downstream analysis as outlined above.

## Data availability

10x multiome data in the study are available in the NCBI Gene Expression Omnibus (GEO) under accession number GSE229169 and for viewing on the WashU Comparative Epigenome Browser data hub: https://epigenome.wustl.edu/BrainComparativeEpigenome/

## ACKNOWLEDGEMENTS

We thank all other members of the Ren and Ecker laboratory for their valuable inputs. This study was supported by NIH Grants U19MH11483 to J.R.E and E.M.C, U19MH114831-04s1 to J.R.E. and B.R., 5U01MH121282 to J.R.E and M.M.B and UM1HG011585 to B.R. J.R.E. is an investigator of the Howard Hughes Medical Institute. Work at the Center for Epigenomics was also supported by the UC San Diego School of Medicine. This publication includes data generated at the UC San Diego IGM Genomics Center utilizing an Illumina NovaSeq 6000 that was purchased with funding from a National Institutes of Health SIG grant (#S10 OD026929).

## AUTHOR CONTRIBUTIONS

Study supervision: B.R., J.R.E. Contribution to data analysis: N.R.Z., E.J.A. W.W., J.Z., S.L., H.L., Y.E.L., W.T. Contribution to data generation: N.R.Z, J.R.N., R.C., A.B., M.M. Contribution to data interpretation: N.R.Z., E.J.A., W.W, B.R., J.R.E., Contribution to writing the manuscript: N.R.Z., E.J.A, B.R., W.W. All authors edited and approved the manuscript.

## Supplementary Information

Supplementary Table 1 Conservation GLS T-statistic for gene expression.

Supplementary Table 2 Pairwise differential gene expression.

Supplementary Table 3 Conservation and divergence annotation of human protein coding gene expression.

Supplementary Table 4 Human ATAC peak summary.

Supplementary Table 5 Macaque ATAC peak summary.

Supplementary Table 6 Marmoset ATAC peak summary.

Supplementary Table 7 Mouse ATAC peak summary.

Supplementary Table 8 Conservation and divergence annotation of human atac-peaks.

Supplementary Table 9 Conservation GLS T-statistics for chromatin accessibility.

Supplementary Table 10 Pairwise differential chromatin accessibility.

Supplementary Table 11 Human DMRs.

Supplementary Table 12 Macaque DMRs.

Supplementary Table 13 Marmoset DMRs.

Supplementary Table 14 Mouse DMRs.

Supplementary Table 15 Conservation of human DMRs.

Supplementary Table 16 Conservation GLS T-statistic for DMRs.

Supplementary Table 17 Human boundaries.

Supplementary Table 18 Macaque boundaries.

Supplementary Table 19 Marmoset boundaries.

Supplementary Table 20 Mouse boundaries.

Supplementary Table 21 Conservation of human boundaries.

Supplementary Table 22 Human loops.

Supplementary Table 23 Macaque loops.

Supplementary Table 24 Marmoset loops.

Supplementary Table 25 Mouse loops.

Supplementary Table 26 Conservation of human loops.

Supplementary Table 27 Human ABC.

Supplementary Table 28 Macaque ABC.

Supplementary Table 29 Marmoset ABC.

Supplementary Table 30 Mouse ABC.

Supplementary Table 31 Conservation of human ABC.

---

**Extended data Fig. 1:**
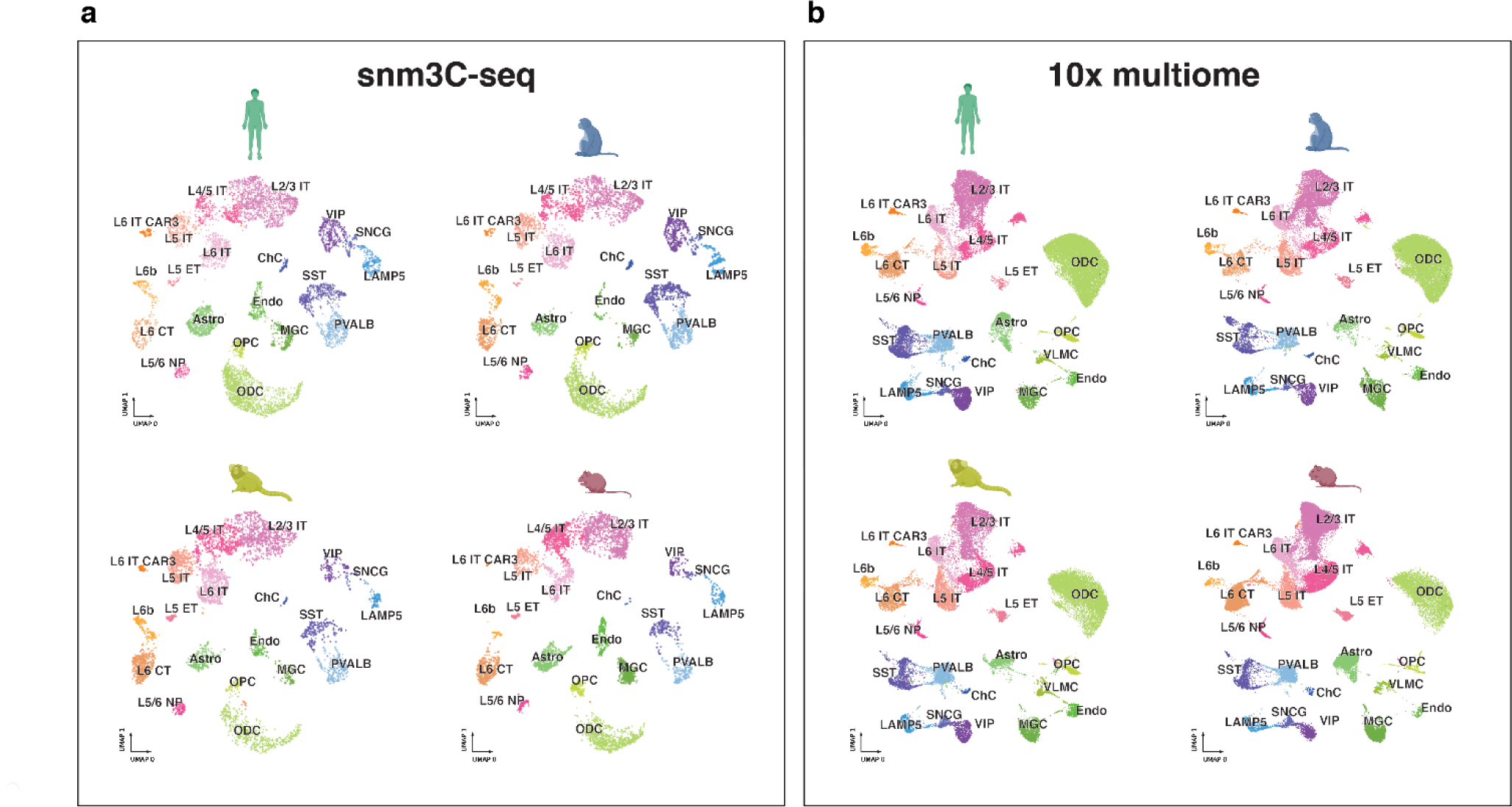
Species separated snm3C-seq and 10x multiome UMAPs. **a.** Uniform manifold approximation and projection (UMAP)^52^ embeddings of snm3C-seq DNA methylation clusters for human, macaque, marmoset, and mouse separately. **b**. same as in **a**, but for 10x multiome RNA.

**Extended data Fig. 2:**
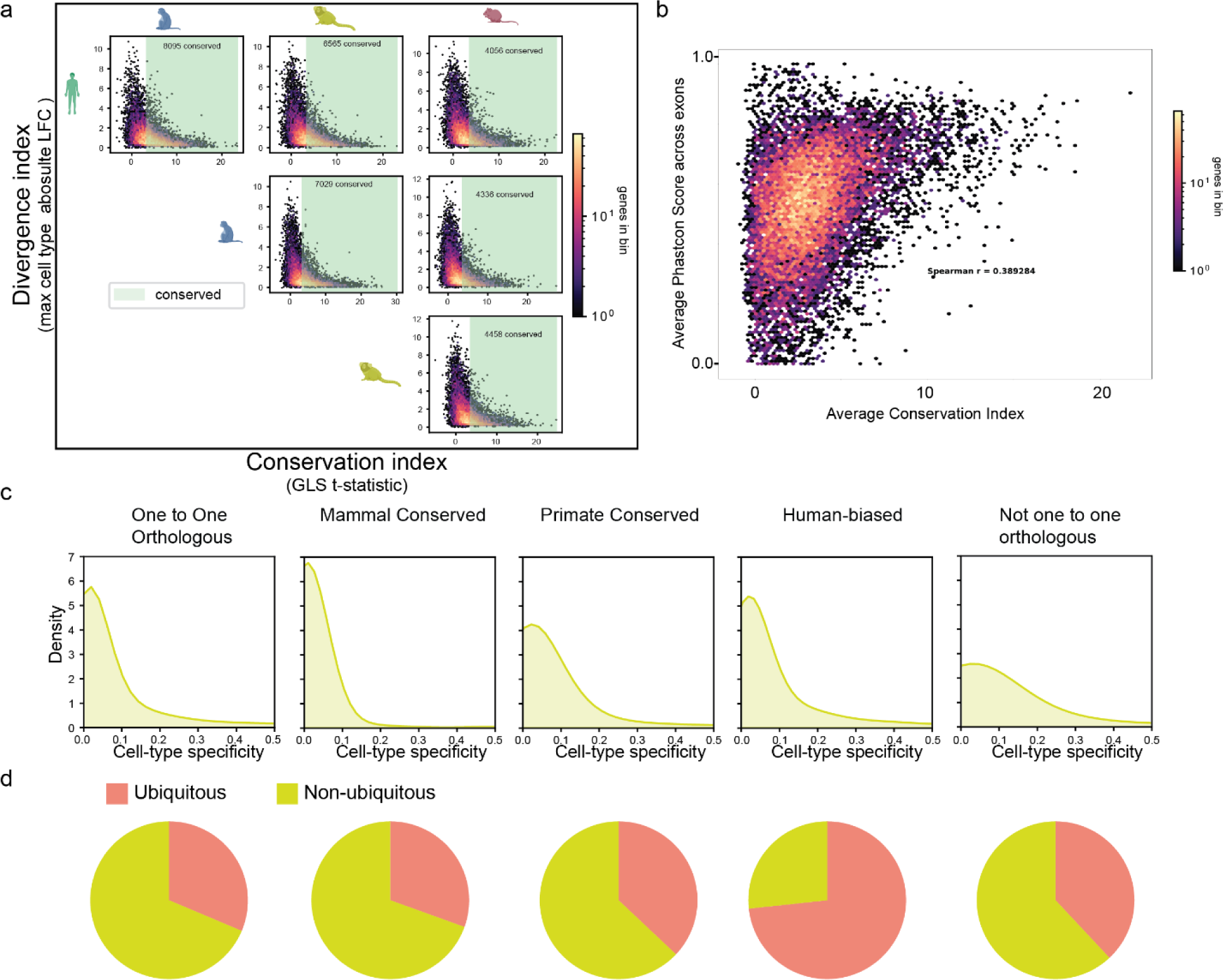
Patterns of gene expression conservation and divergence. **a.** Pairwise divergence vs conservation index of gene expression for each species pair **b.** Correspondence of gene expression conservation to average phastcons score of exons. **c.** Entropy-based cell type specificity across human cell types. **d.** Proportion of ubiquitously expressed genes by category.

**Extended data Fig. 3:**
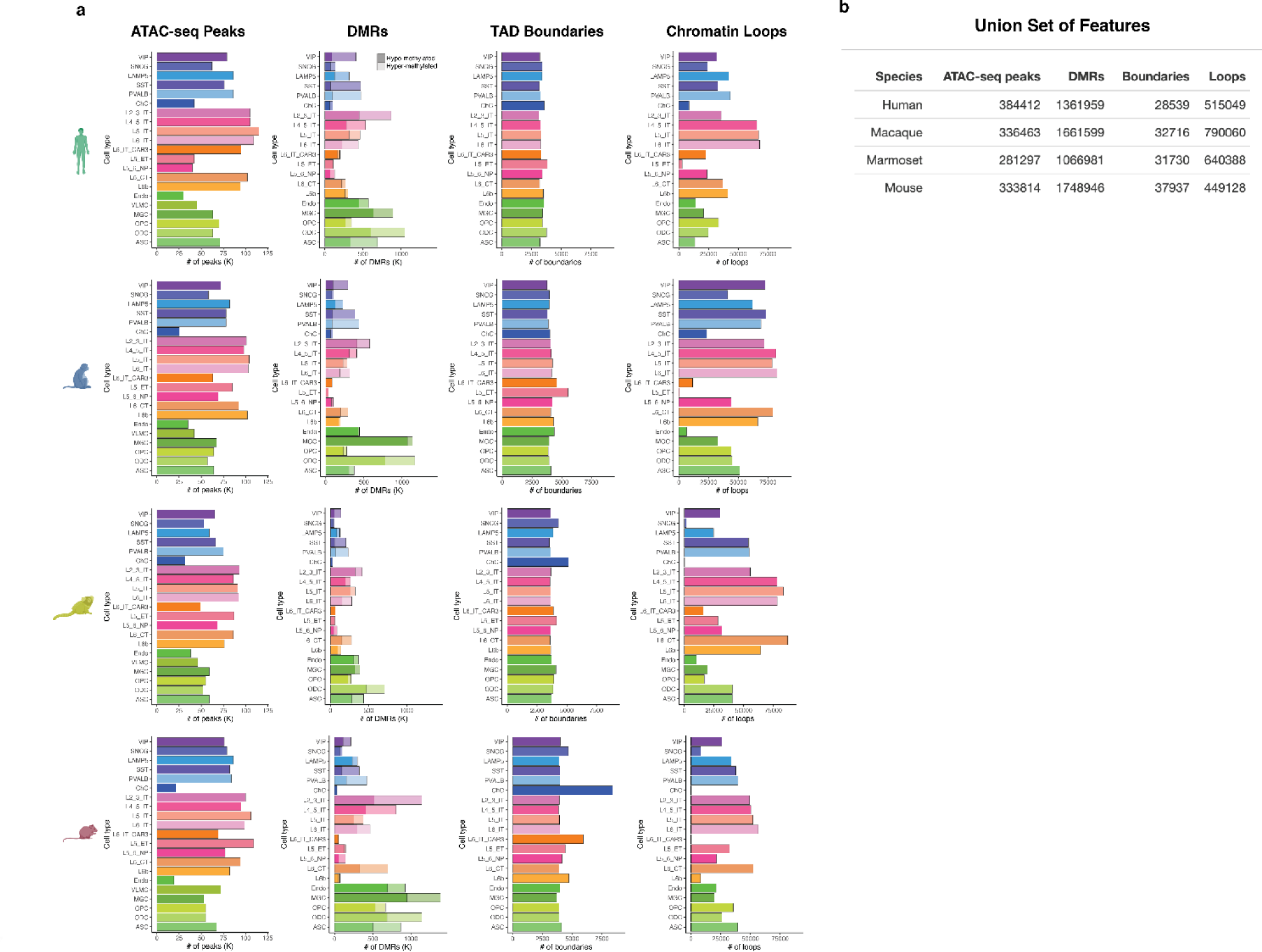
Numbers of features identified in each species and cell type. **a.** Number of indicated features (ATAC-seq peaks, DMRs, TAD boundaries, or chromatin loops) identified for each cell type for each species. **b**. Numbers of unique features found in each species, i.e. union set of features.

**Extended data Fig. 4:**
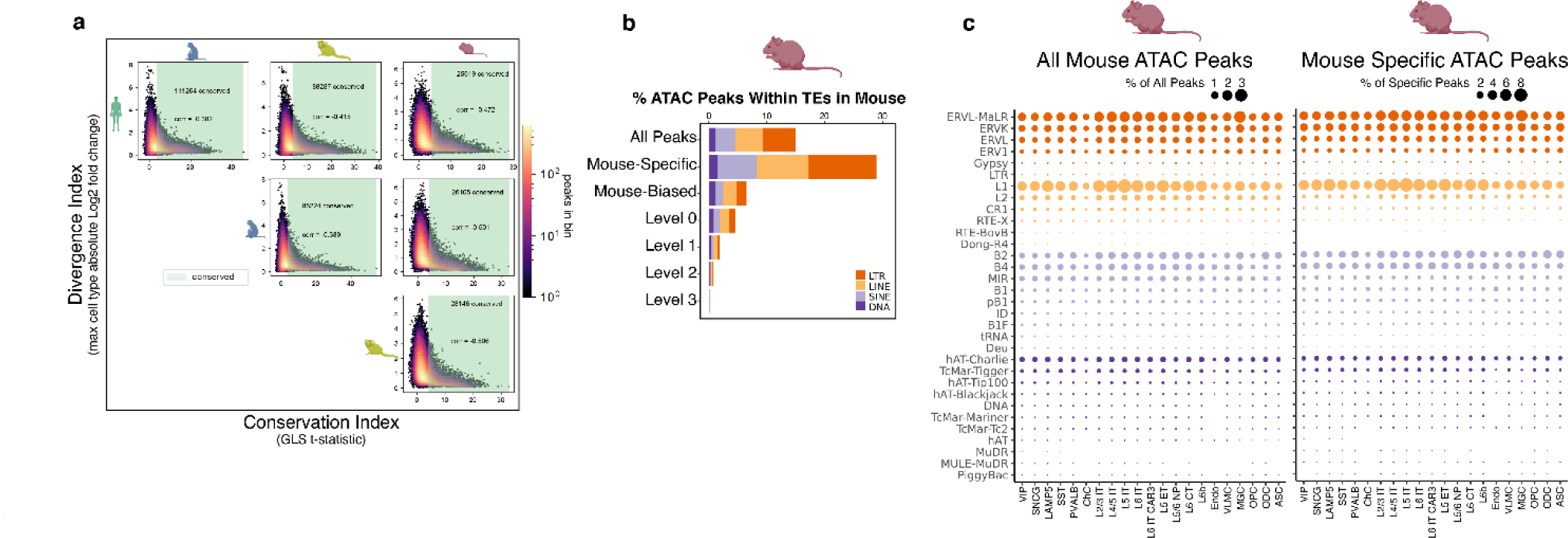
Comparative chromatin accessibility and enrichment in mouse TEs. **a.** Pairwise divergence vs conservation index of ATAC-seq peaks for each species pair. **b.** Stacked bar plots showing percentage of mouse cCREs in TEs for different conservation groups. **c**, Dot plots showing the percentage of all (left) or mouse-specific (right) cCREs in different subclasses of TEs for each cell type.

**Extended data Fig. 5:**
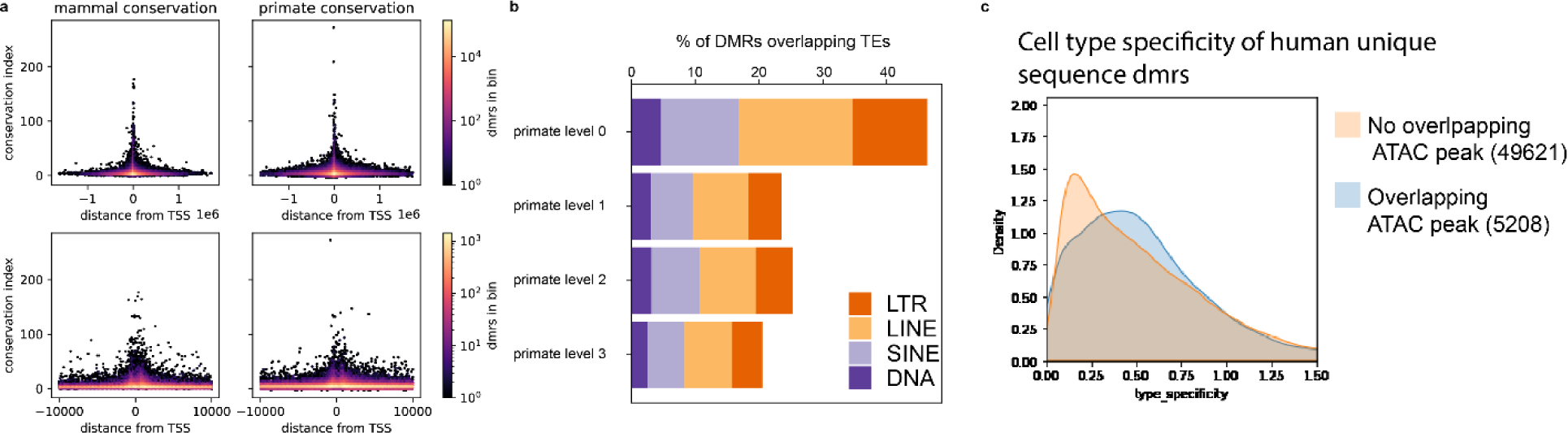
Patterns of DNA methylation conservation. **a.** Correspondence of distance from nearest TSS to DMR conservation level. **b.** Proportion of TEs in different levels of primate conserved DMRs. **c.** Differences in cell type specificity between human-specific DMR sequences which overlap peaks, and human-specific DMR sequences which don’t.

**Extended data Fig. 6:**
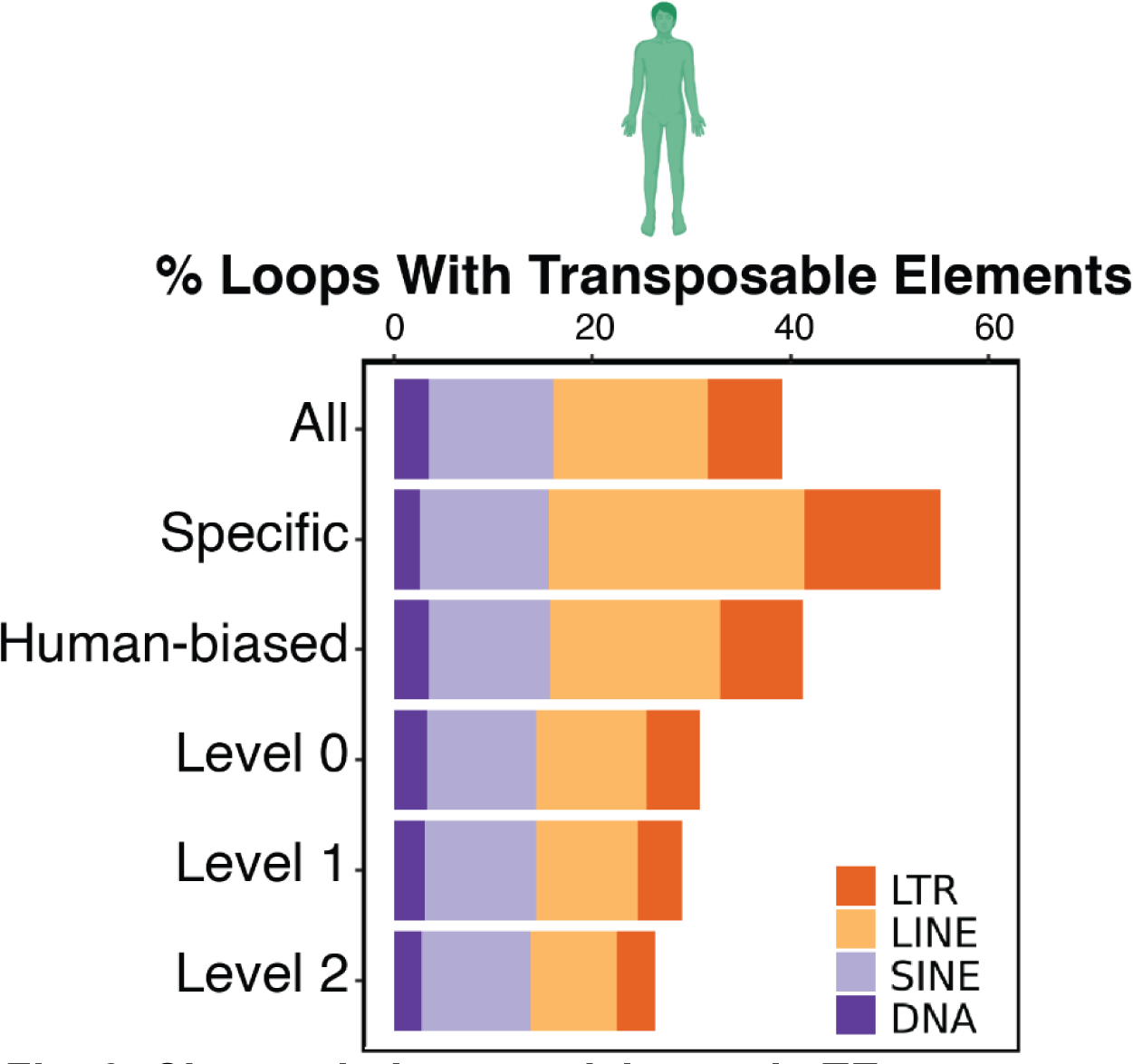
Chromatin Loop enrichment in TEs. **a.** Stacked bar plots showing percentage of human chromatin loops overlapping TEs for different conservation groups.

**Extended data Fig. 7:**
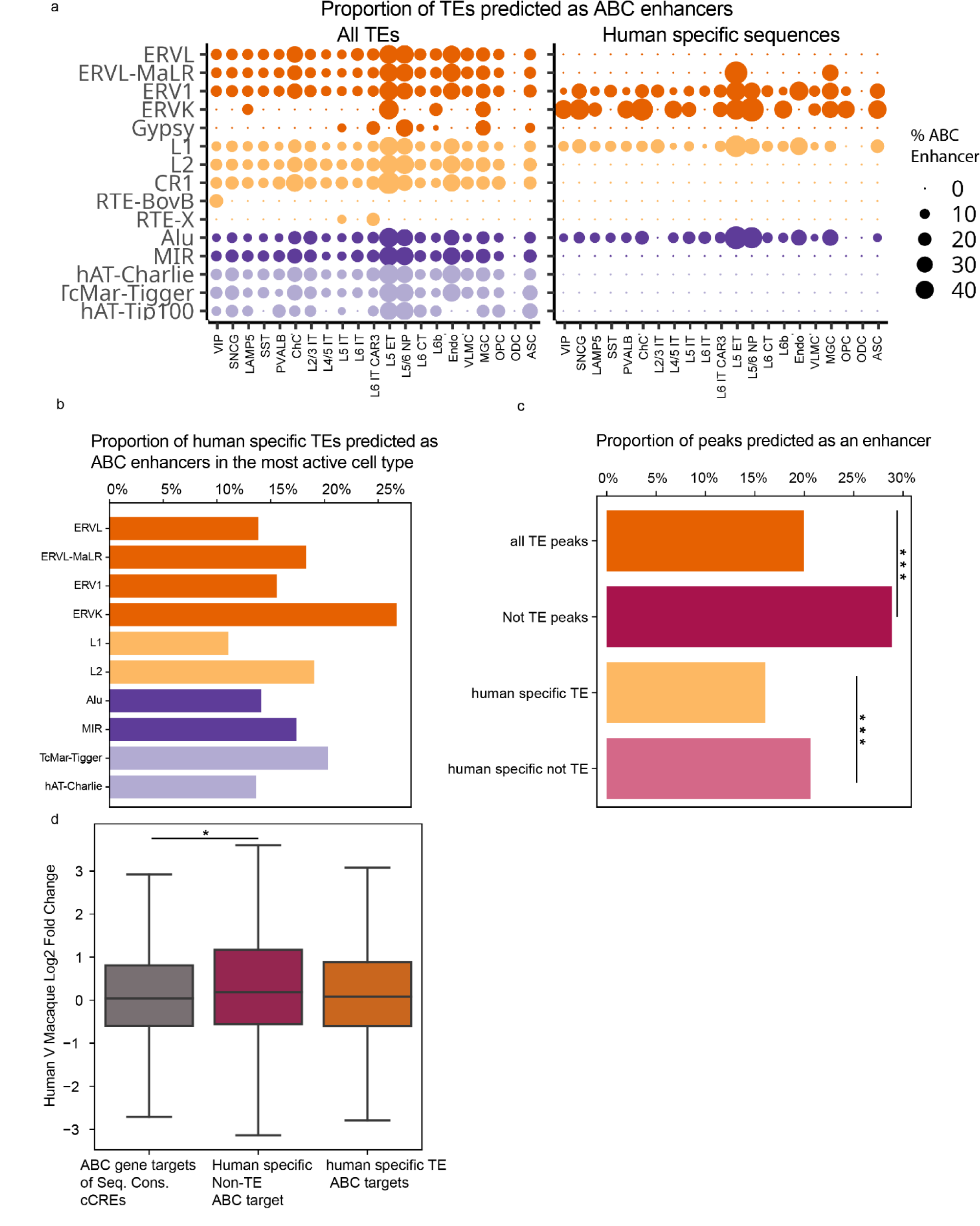
Cell type specificity of predicted transposable element enhancer function. **a.** Proportion of TEs predicted by ABC to be enhancers (ABC score ≥ 0.02) in each cell type by transposable element class. **b.** Proportion of most common transposable element classes predicted to be ABC enhancers (ABC score ≥ 0.02) in their most accessible cell type. **c.** Proportion of ABC predicted enhancer peaks between TE and non-TE peaks, as well as human-specific peaks (*** p < 1e-3, Chi-squared test). **d.** Human vs macaque log2 fold change for ABC targets of human-specific sequences (* p < 0.05, Mann-Whitney U test).

**Extended data Fig. 8:**
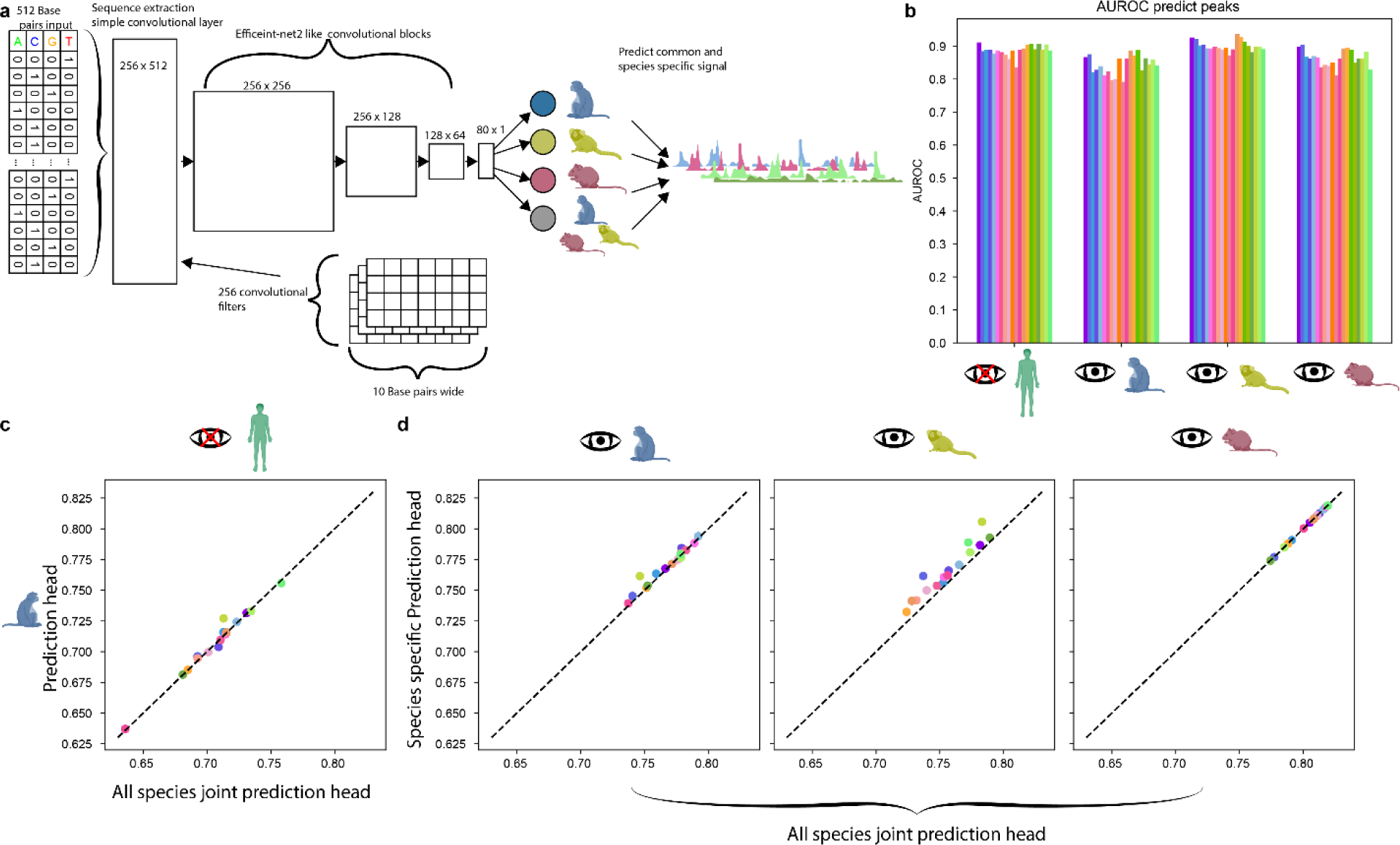
Species specificity of open chromatin deep learning. **a.** A schematic highlighting further details of neural network implementation. **b.** AUROC for peak regions based upon predicted signals for each species. **c.** Predicted accuracy within cell types using the macaque-specific prediction output for human test set, vs. using all species predicted output. **d.** Comparison of correlation across bins within a cell type between the species-specific predictive head, and all species predictive head.

